# Learning to represent continuous variables in heterogeneous neural networks

**DOI:** 10.1101/2021.06.01.446635

**Authors:** Ran Darshan, Alexander Rivkind

## Abstract

Animals must monitor continuous variables such as position or head direction. Manifold attractor networks—which enable a continuum of persistent neuronal states—provide a key framework to explain this monitoring ability. Neural networks with symmetric synaptic connectivity dominate this framework, but are inconsistent with the diverse synaptic connectivity and neuronal representations observed in experiments. Here, we developed a theory for manifold attractors in trained neural networks, which approximate a continuum of persistent states, without assuming unrealistic symmetry. We exploit the theory to predict how asymmetries in the representation and heterogeneity in the connectivity affect the formation of the manifold via training, shape network response to stimulus, and govern mechanisms that possibly lead to destabilization of the manifold. Our work suggests that the functional properties of manifold attractors in the brain can be inferred from the overlooked asymmetries in connectivity and in the low-dimensional representation of the encoded variable.

## 1 Introduction

Stimulus-specific neuronal activity persists even without the stimulus [Wang, 2001, Constantinidis et al., 2018]. Such persistent states of neuronal circuits were hypothesized to be sustained by recurrent synaptic connections [Hebb, 1949, Durstewitz et al., 2000] and are called neural *attractors* [Hopfield, 1982, Amit, 1992]. In a wide variety of brain systems the variables that are represented by such persistent neuronal activity are continuously valued [Funahashi et al., 1993, Romo et al., 1999]. In particular, neurons in the navigation system represent continua of animals directional heading [Taube et al., 1990, Seelig and Jayaraman, 2015], speed [Kropff et al., 2015] and locations [O’Keefe and Dostrovsky, 1971, Hafting et al., 2005], while neuronal activity in prefrontal and posterior parietal neocortices correlates with stimulus orientation [Christophel et al., 2017] and its spatial location [Funahashi et al., 1993].

A common framework to study such continuous internal representations is the theory of computations by *manifold* attractor networks [Amari, 1977, Ben-Yishai et al., 1995, Seung, 1996, Burak and Fiete, 2009, Wimmer et al., 2014, Hansel and Mato, 2013, Chaudhuri et al., 2019, Gardner et al., 2021]. In these networks, the variable of interest is represented as a point in the space of neural activity, with the continuum of values forming a low-dimensional manifold of attractor states in the high-dimensional space of neural firing rates. Neural dynamics converge toward the manifold attractor, and are thus robust to perturbations that could kick the state away from it. On the other hand, due to the continuum of stable states, stability is *marginal* [Ben-Yishai et al., 1995] along the manifold: a perturbation in this direction does not face either converging nor repelling forces [Durstewitz et al., 2000, Chaudhuri and Fiete, 2016]. These dynamical features support the computational capabilities of such networks to track external features and do path integration [Hulse and Jayaraman, 2020], or memorize continuous features through persistent activity [Compte et al., 2000].

Most models of manifold attractors strongly rely on symmetry assumptions [Amari, 1977, Brody et al., 2003, Machens and Brody, 2008, McNaughton et al., 2006, Burak and Fiete, 2009, Chaudhuri and Fiete, 2016, Mastrogiuseppe and Ostojic, 2018, Beiran et al., 2020]. For example, in models of the head-direction system [Zhang, 1996], or in representations in the primary visual [Ben-Yishai et al., 1995] and prefrontal [Compte et al., 2000] cortices, the connectivity is constructed according to a rotation symmetry principle, in which recurrent interactions depend only on the distance between the preferred direction of the neurons. As a result, the connectivity profile is the same for all neurons up to a rotation of the angular feature (Fig. 1Ai-ii, see also [Mastrogiuseppe and Ostojic, 2018] for an extension for these connectivity rules). Under some general conditions on the recurrent interactions, such as short-range excitation and long-range inhibition, a manifold of attractors appears (Fig. 1Aiii). We call these classical models which are based on a symmetry assumption, *symmetric-connectome* attractor networks.

**Figure 1:**
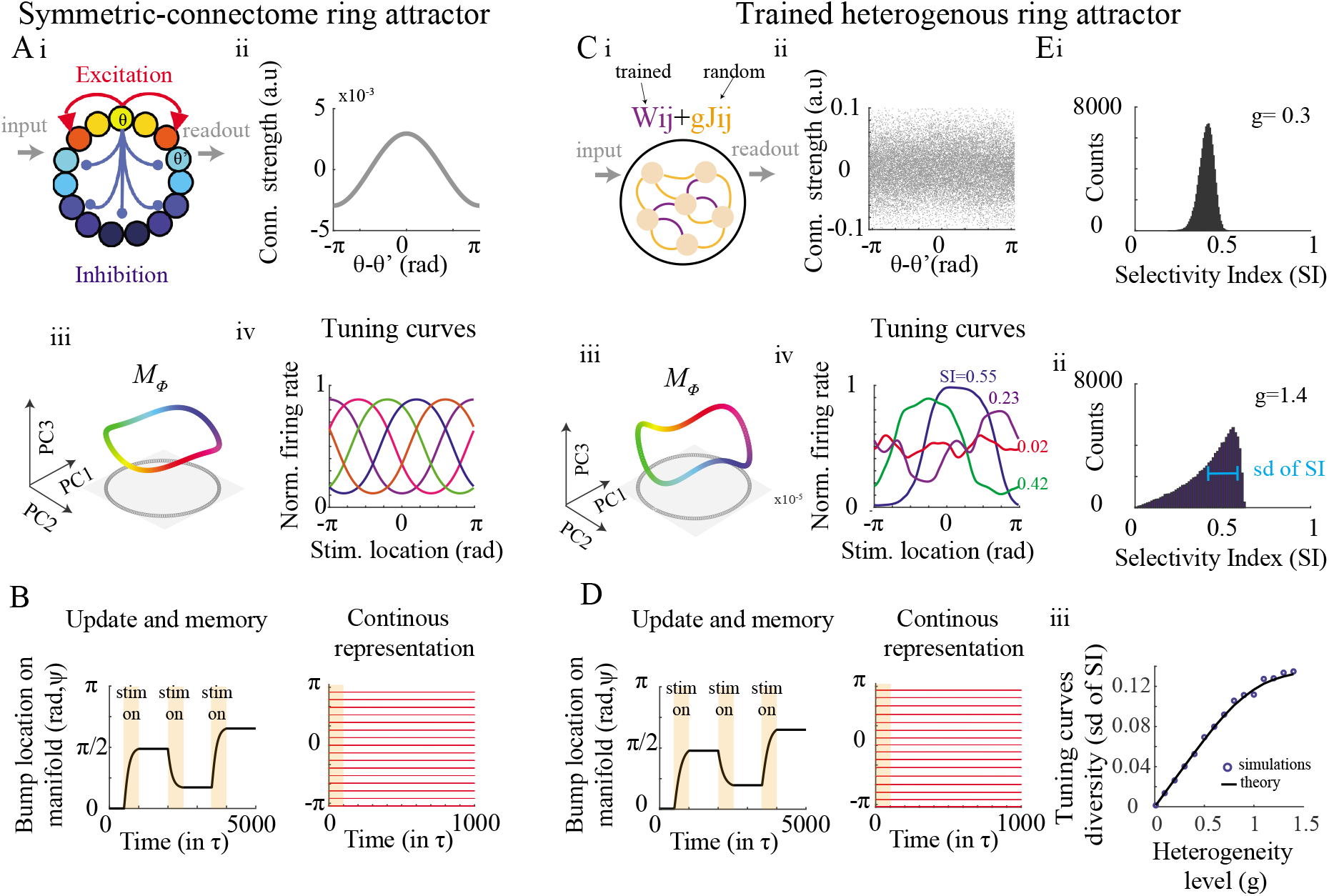
The symmetric-connectome ring attractor network and heterogeneous trained ring attracor network. **A,B.** Symmetric-connectome ring model. **Ai.** Cartoon of a ring network architecture. Neurons are aligned on a ring, in which the connectivity depends solely on the difference between the neuronal preferred directions, denoted by *θ*. **ii.** Rotation symmetry in the connectivity: connectivity strengths coincide for all neurons in the network when plotted against the difference between the neuronal preferred directions. **iii.** The manifold attractor, denoted by *M_ϕ_*, projected on the leading PCs of the neural representation (i.e. the tuning curves). Color indicates the decoded representation of the continuous feature (the angle of the decoder in Eq.(3)). Note that for a ring manifold the projection on the first two PCs exhibits a circular shape (gray). **iv.** Tuning curves of example neurons in the symmetric-connectome model are all identical, up to a symmetry for rotations. **B.** Left: Bump’s location on the manifold (in radians) vs. time (in units of membrane time-constant). The internal representation of the feature (a ‘bump’ of activity) evolves when external input is applied (orange areas) and persists for a long time without input (white areas). Right: Continuous internal representation of the feature. Red lines: position of the bump following initialization at that locations using the external input (in orange). The state persists without inputs. **C-D.** Same as (A-B), but for a *trained* heterogeneous ring. Connectivity consists of random heterogeneous component, *gJ_ij_* (orange connections) superimposed with the structured component, *Wij* (purple); only the latter component is affected by training. Contrary to the case in panel **A**, connectivity is not fully determined by difference in preferred directions **(ii)** and tuning curves do exhibit diversity **(iv)**. **E.** Diversity in tuning curves increases with the amount of synaptic heterogeneity. **Ei-ii.** Examples of distribution of selectivity index (SI), defined as the first Fourier component of the tuning curve (see Methods) for networks that were trained with different amounts of synaptic heterogeneity. See also examples in Civ. **Eiii** Diversity in tuning curves (SD of SI, see Eii) increases with the heterogeneity level. Theory in Eq.(54)

The symmetry assumption is highly unlikely in real biological systems, in which heterogeneity in synaptic connections is abundant [Braitenberg and Schüz, 2013]. Yet, in these classical models, any deviation from perfect symmetry shatters the continuous attractor into a few isolated attractors, leading to a fast deterioration of the computational capabilities of the network, such as the loss of persistent representation [Zhang, 1996, Tsodyks and Sejnowski, 1995, Renart et al., 2003, Itskov et al., 2011] or imperfect path integration [Burak and Fiete, 2009]. Furthermore, as a direct outcome of this symmetry in connectivity, the activity profile of neurons in symmetric-connectome models are identical. This is in sharp contrast to neurons in reality that can show many degrees of diversity [Barak et al., 2013, Finkelstein et al., 2015, Chaudhuri et al., 2019, Fisher et al., 2019].

Beyond these constraints on the synaptic connections and neuronal tuning profile, symmetric-connectome models imply, by their construction, a perfect *geometry* of the high-dimensional neuronal representation along the manifold. In particular, a one-dimensional manifold will have a perfect circular shape when projected to the leading principal components of neural activity (Fig. 1Aiii) and average activity, which is independent of the location on the manifold. Such a perfection of geometry in the brain is not supported by experimental evidence: while recent studies suggest that neuronal representations of continuous features might exhibit the topology of a ring [Chaudhuri et al., 2019, Rubin et al., 2019] or a torus [Gardner et al., 2021], in agreement with the manifold attractor hypothesis, there is no evidence for a perfect geometrical symmetry in these representations. Symmetric-connectome attractor models are thus inconsistent with synaptic heterogeneity and diverse tuning profiles of neurons and cannot support imperfect geometries in neural manifolds.

To loosen these idealistic assumptions, we considered a manifold that emerges as a result of learning rather than via a pre-engineered connectivity. Indeed, trained on a wide variety of tasks, ranging from integration of evidence [Mante et al., 2013], to path integration [Sorscher et al., 2020, Cueva et al., 2019] and natural language processing [Maheswaranathan et al., 2019], recurrent neural networks (RNNs) exhibit manifold attractor dynamics. In these unconstrained settings, when symmetry in the connectivity is not imposed and models with synaptic heterogeneity are trained, neither the connectivity nor the geometry of the representation is expected to exhibit a perfect symmetry. However, we are lacking a theoretical understanding of how manifold attractors emerge in these models or how the neuronal representation in the trained RNNs shapes their dynamical properties and computational power.

How can a continuum of persistent states co-exists with the synaptic heterogeneity and diversity of neuronal representation observed in experiments? What is the effect of such diversity and of the manifold’s geometry on the dynamics along the manifold and in its vicinity? We present a minimal and solvable model of a trained recurrent network that we analyzed analytically in the large network limit using mean-field approach, and in which we relax the symmetry assumption in both synaptic connectivity and manifold geometry. We show that in such networks, a continuous representation emerges from a small set of stimuli used for training. As in real biological systems, tuning curves and connectivity patterns in the model are heterogeneous. Furthermore, our framework encompasses biological manifolds with non-symmetric geometry, in sharp contrast to existing theories that can only deal with symmetric manifolds. Finally, our theory connects synaptic heterogeneity and imperfect geometry of the internal representation to the computational properties of the attractor network, such as its robustness to perturbation and its response to external stimulus. Our work thus shows that diversity in synaptic connectivity and imperfections in neural representation can coexist with the hypothesis of neural manifold attractors, obviating the need to rely on symmetry principles.

## 2 Results

We studied networks consisting of *N* firing rate units:

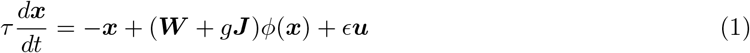

where *x_i_* is the total input into the *i*-th neuron, and the recurrent connectivity is decomposed into a structured connectivity matrix, ***W***, and a random matrix, ***J***. The strength of the external input, ***u***, is controlled by a parameter *ϵ*, and the level of synaptic heterogeneity in the recurrent network is controlled by a parameter *g*. We used a sigmoidal transfer function, *ϕ*(*x*), but it is possible to generalize our theory to other transfer functions.

We will use the term symmetric-connectome network models to refer to the case without heterogeneity (*g* = 0) and where the structured recurrent component is constructed according to symmetry principles, instead of via training (Fig. 1A). Indeed, many computational and theoretical studies use these types of network models to explain the emergence of continuous representations in visual cortex, continuous and persistent activity in prefrontal cortex, and persistent activity and path integration in the navigation system. In such models, a continuous periodic feature, ψ, is encoded in the network, and the recurrent connectivity is invariant to rotations (Fig. 1Ai-ii). Without external inputs, all states are equally stable and form a manifold of attractors, i.e., activity of the neurons lies on a one-dimensional manifold (Fig. 1Aiii). Each state is a packet, or a ‘bump’ of localized activity. With external cues, the symmetry is broken, and an appropriate state is selected from the continuum of states on the manifold. Figure 1B depicts both update of the memorized feature when the stimulus is present and periods of persistent activity when it is absent.

In what follows, we show that while symmetric-connectome network models [Ben-Yishai et al., 1995, Burak and Fiete, 2009, Beiran et al., 2020, Mastrogiuseppe and Ostojic, 2018], such as the one depicted in Fig. 1A, are a possible implementation for computations based on manifold attractors, they are by far not the only ones.

### 2.1 Trained manifold attractors

In contrast to symmetric-connectome models, we assume that the structured component is *trained* rather than set *a priori*. We then write the structured recurrent connectivity in the following form:

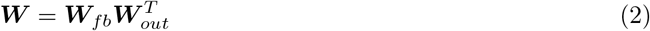

which allows us to interpret this recurrent component as a mapping from the neural activity into a two dimensional (2D) representation which we denote by ***z***, via ***W**_out_*:

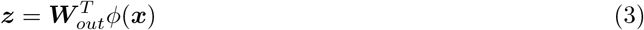

and ***W**_fb_* projects it back to the neurons [Jaeger, 2001, Sussillo and Abbott, 2009]. Specifically, we train the structured connectivity such that the output, ***z***, lies on a desired, predefined, manifold. The continuous feature *ψ* is then read out from the network through the angle of the 2D vector ***z***. To clarify notations, we distinguish the manifold of internal states *ϕ* ∈ *M_ϕ_* (Fig1A,Ciii), which is embedded in the N-dimensional neuronal state, from the pre-defined trained manifold projected in 2D, 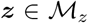 (e.g. see Fig. 1Aiii, Fig. 3 and Methods).

Figure 1C shows an example of such a network that was trained with synaptic heterogeneity (*g* > 0) and without relying on a symmetric-connectome. Specifically, we trained the network to produce a ring manifold, i.e., requiring a circle of radius A in the *decoder plane*: ***z***(*ψ*) = *A*[cos(*ψ*), sin(*ψ*)]. As a result of training, the activity in the network persists for a long time, and the network memorizes the angular feature, until the representation gets updated through an external stimulus (Fig. 1D). This is in contrast to models with pre-defined symmetric-connectome [Ben-Yishai et al., 1995, Mastrogiuseppe and Ostojic, 2018], where both memorization and update capabilities are impaired by heterogeneous connectivity (Fig. S1).

The neuronal representation in the trained network is dramatically different from its symmetric-connectome counterpart. Due to the presence of synaptic heterogeneity, not all states are identical; both the population activity profile (the ‘bump’), and the tuning curves of neurons in the model are highly heterogeneous. Figure 1Civ and Fig. S2 exemplifies such highly heterogeneous tuning curves. To quantify this property, we solved the steady state of Eq.(1) and used it to derive the statistics of the tuning curves in the model (see Methods). The diversity in tuning curves increases monotonically with the amount of synaptic heterogeneity in the network (Fig. 1E). These analytical calculations are in good agreement with our simulations (compare solid line with circles).

Did learning result in a structured component ***W*** that merely compensates for the heterogeneous component *g**J***, such that it restores the rotational symmetry of the synaptic connectivity? We find that this is not the case. The recurrent connectivity is not solely a function of the distance in preferred directions, as is the case for the symmetric-connectome models (compare Fig1Aii with Fig1Cii). This is true also when considering only the structured component of the recurrent interactions, which is not symmetric (Fig. S2). Indeed, it is only after considerable averaging of synaptic inputs across neurons that we can observe that the connectivity profile depends on the distance in neuronal preferred directions (Fig. S2). Finally, symmetry was restored only in a specific case in which we trained the network without any heterogeneity (*g* = 0).

Figure 1C shows the existence of a manifold attractor without relying on symmetry in the synaptic interactions or neuronal tuning curves. Yet, in this case due to the circular shape of the manifold in the 2D plane (***z***(*ψ*)), the second-order statistics of the neuronal representation are invariant for rotations, and the average population activity is the same at each point on the manifold (black line in Fig. 2A). Consequently, the principal components (PCs) of the neural representation are the spatial Fourier modes, and the projection of the manifold on the leading PCs features a circular shape (note the gray circle in Fig. 1Ciii). However, symmetry in the second order statistics is not necessary for the emergence of manifolds attractors. Indeed, Figure 2 shows an example of a trained manifold that is not circular. In this example, the average population activity varies along the manifold (gray lines in Fig. 2A), the distribution of preferred direction can be non-uniform (Fig. 2F), the second-order statistics of the representation show no rotation symmetry (Fig. 2G), and the PCs are not pure Fourier modes (not shown). However, similarly to the ring manifold, these networks can memorize and update the representation of a continuous feature based on manifold attractor dynamics (Fig. 3).

**Figure 2:**
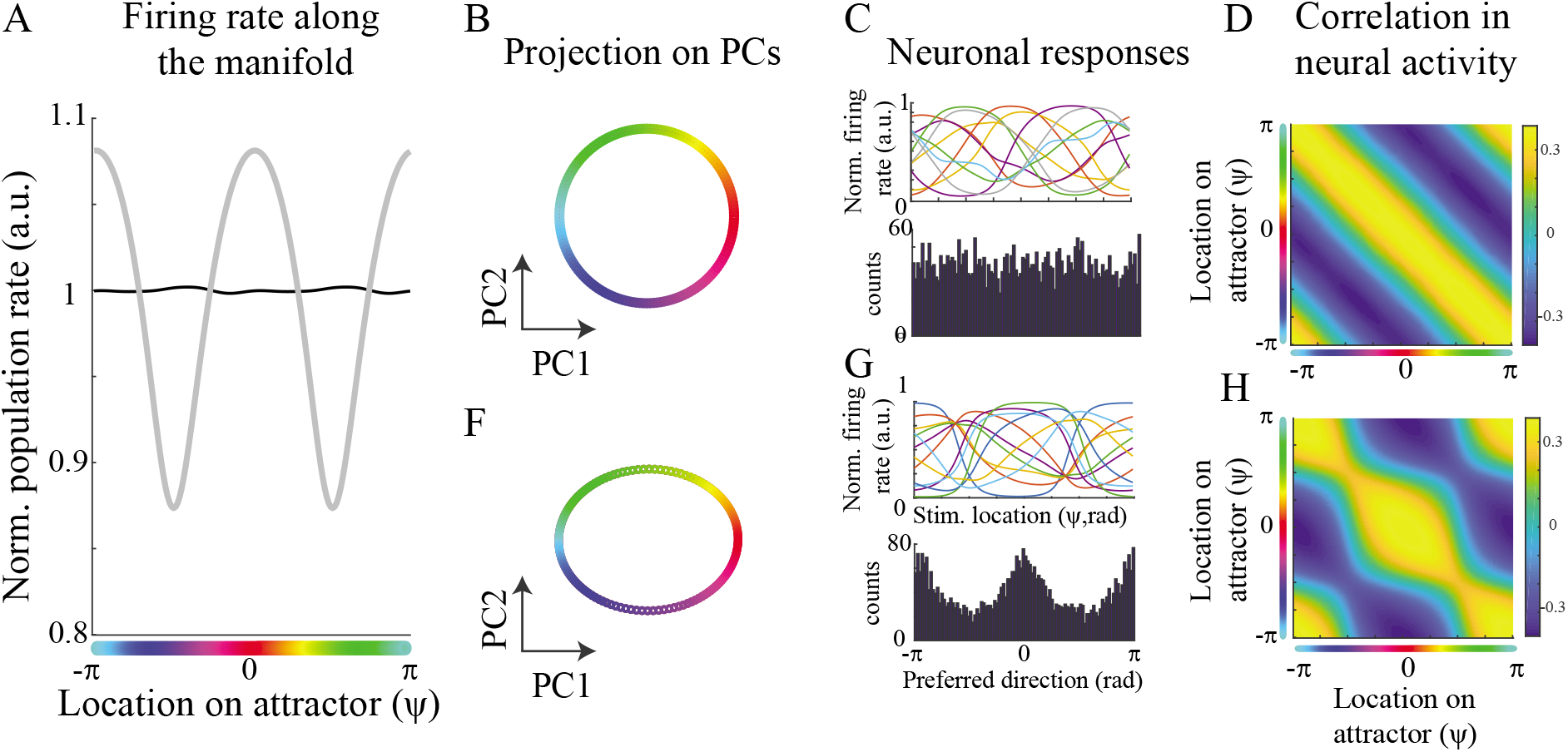
Neural representations of manifold attractors. **A**. Normalized population activity along the manifold (Eq.(10)) for a ring manifold (black, B) and the ellipse manifold in E (gray). In a ring manifold the total population activity is constant at each location of the manifold (up to small fluctuations that arise in small networks). **B.** Projection of the neural representation on the top two leading principal components for a ring manifold (see parameterization of the trained manifold in Methods). The represented feature, ψ, is color coded. **C.** Top: Tuning curves of neurons in the trained network. Bottom: distribution of preferred directions in the network. **D.** Correlation across population activity at different locations on the manifold, *C*(*ψ, ψ′*) = 〈*ϕ*(***x***(*ψ*))*ϕ*(***x***(*ψ′*))〉. For a perfect ring geometry the correlation function exhibits a rotation symmetry (*C*(*ψ,ψ′*) = *C*(*ψ* – *ψ′*), i.e. matrix is circulant). **E-F**. Same as (B)-(D) but for an ellipse manifold in which there is no rotational symmetry in the representation

**Figure 3:**
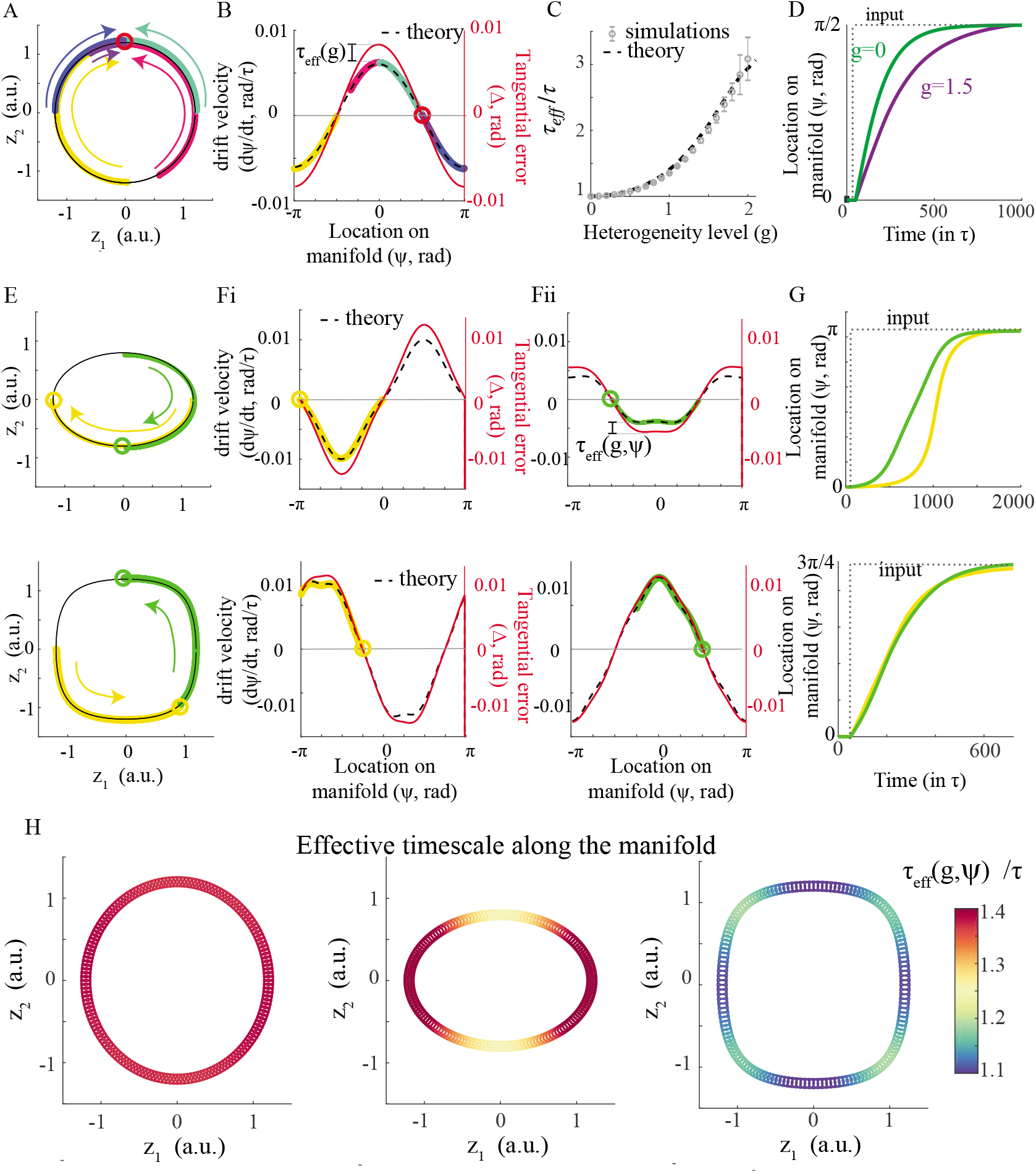
Translation on manifold attractors in response to external inputs A-D. Trained ring manifold. **A.** Projection of neural activity on the decoder (***z*** plane) for a trained ring manifold with external input at *ψ* = *π*/2 (red circle). All five trajectories (different colors), starting from random locations on the manifold, converge to *ψ* = *π*/2. Trajectories were slightly adjusted to not overlap for illustration purposes. **B.** Drift velocity of the trajectories in (A) (same color code) against location on the manifold. Red circle: angle of the external input. Red curve: Tangential error. Dashed line: Theory (Eqs.(4), (5)). The difference between the tangential error and the drift velocity is a simple scaling by *τ_eff_*. **C.** The effective timescale increases with the amount of heterogeneity in trained ring manifolds. Dashed line: theory from Eq.(5). Gray: S.E.M for 10 network realizations. **D.** Bump’s location vs. time for networks that were trained with different amounts of synaptic heterogeneity. The external input is presented at *t* = 50*τ* at *ψ* = *π*/2. The network’s internal representation rotates toward *π*/2 with a velocity that depends on the amount of synaptic heterogeneity. **E-G.** Top: Trained ellipse manifold from Fig. 2E-G. Bottom: another example of a trained manifold (see Methods for parameters). **E.** Green trajectory: the bump’s location drifts along the manifold toward the input at *ψ* = –*π*/2 (green circle). Yellow trajectory: drift toward the input, which is now at *ψ* = *π* (yellow circle). **F.** Same as (B) but for the two trajectories in (E). Both the predicted and the actual drift velocities are distorted with respect to tangential error. This is because the effective timescale now depends on the location along the manifold. **G.** Bump location vs. time. Same trajectories as in (E). **H.** Effective timescale along the manifold for the three examples in (A-G). Note the increase in timescale along the corners of the 2D projection *z*, which correspond to high firing activity.

To conclude, we obtain a family of manifold attractor networks that lack obvious symmetry by introducing an appropriately structured low-rank component of the form of Eq.(2) on top of synaptic heterogeneity. The hand-crafted symmetric connectome networks that have been theorized to underlie computations in the brain are only one specific choice of the structured connectivity ***W***, and without the heterogeneous part (*g* = 0), while other, learnable, solutions do not rely on any clear symmetry property.

### 2.2 Dynamics along the manifold

How does the absence of symmetry affect the properties of such learnable manifold attractors, and what are their functional implications? Can heterogeneous networks with asymmetry in the neuronal representation encode continuous variables, such as head direction, while interacting with external stimuli and remaining robust to perturbations? We find that these functional properties are affected when symmetry is lacking.

As a result of the difference in the timescale of the dynamics along the on- and off-manifold directions, they can be analyzed separately. The on-manifold direction is of interest for input driven dynamics and drift, while off-manifold direction accounts for robustness to perturbations.

We first focus on the on-manifold direction. In case that the state is not perfectly persistent due to an external input (*ϵ* > 0, Figs.S3F-H and Fig. 3), an imperfect training (Fig. 4), or changes in the connectivity (Fig. S1), the trajectories quickly converge to the manifold, and the dynamics along the manifold are governed by the following rule:

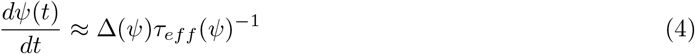

which dissects the speed of motion along the manifold into two factors with a simple interpretation: *τ_eff_* represents the change in the time constant that governs the dynamics along the manifold, while Δ reflects the inconsistency between the current neural state and a perfectly persistent one. This inconsistency, which we term the *tangential error*, is quantified via an auxiliary setting, which we name a *recurrent autoencoder* (RAE). In such a setting, illustrated in Figure S3A-B, and explained in detail in Methods (Section 6.3), we test if a point on the manifold or in its vicinity is a persistent state of the dynamics.

**Figure 4:**
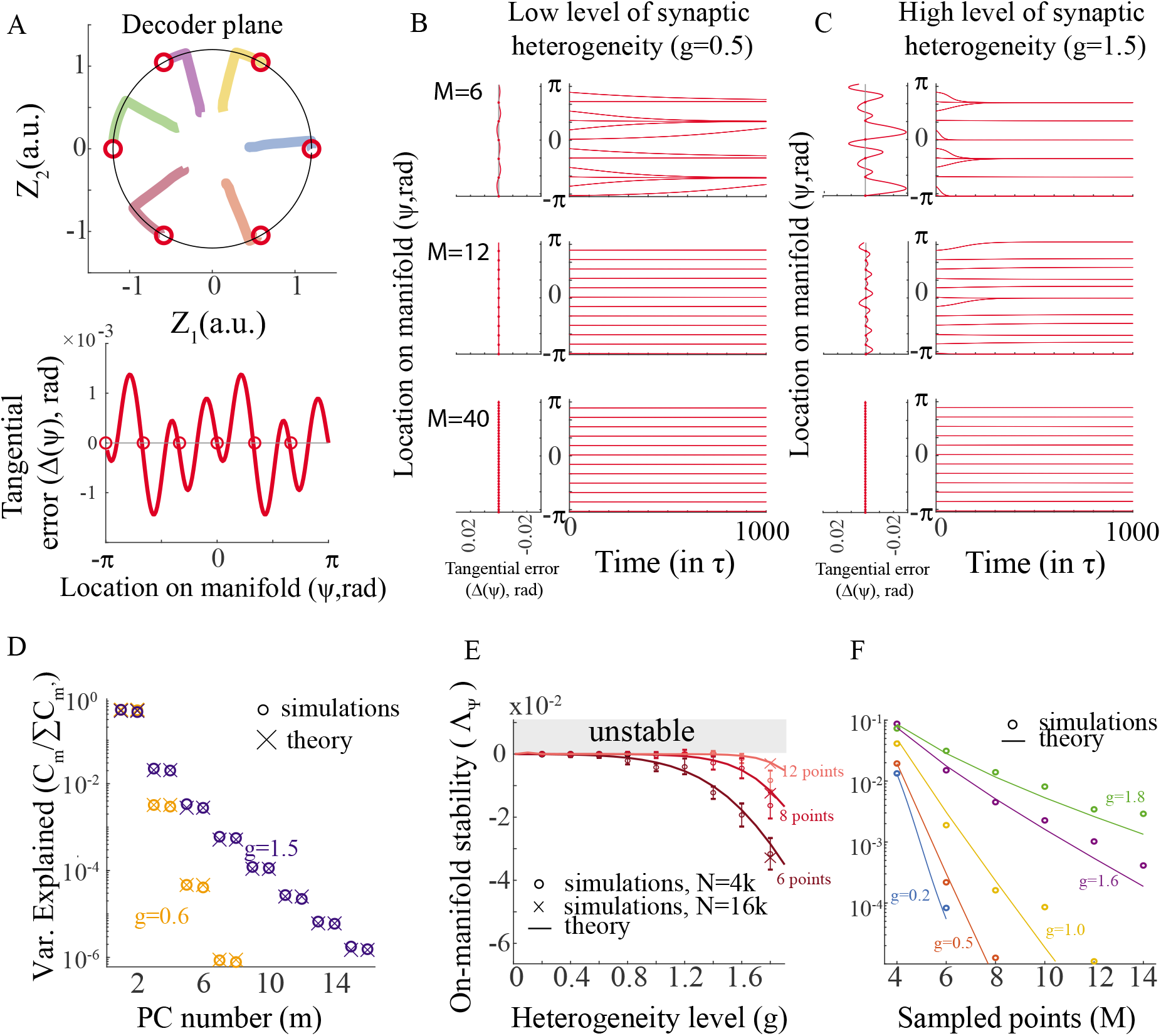
Manifold continuity quickly emerges in trained heterogeneous ring network. **A.** Top: Decoder view of a trained ring manifold when training M=6 equally distributed points on the manifold. Circles: trained points. Colored lines: trajectories starting from different initial conditions quickly converge toward the manifold and then drift slowly to one of the trained points. Bottom: The tangential error of the RAE for the trained network. **B.** Left: The tangential error against the the bump location. Right: Bump location when starting from different points on the manifold as function of time. Synaptic heterogeneity in the network is *g* = 0.5. Top: training the network by sampling M=6 points of the manifold. Middle: same but with M=12. Bottom M=40. **C.** Same as (B), but with g=1.5. **D.** Variance explained (VE) against the principal components number for low (orange) and high (blue) levels of synaptic heterogeneity. Circles: VE as calculated from a simulated network. Crosses: Theoretical prediction (Eq.(46)). **E.** Stability along the manifold direction against the heterogeneity level for M=6, 8 and 12 sample points. Theory: Eq.(6). Note the scale of the y-axis. **F.** Same as (E) but against the number of sampled points. Note the exponential decay to marginal stability (zero eigenvalue), and hence continuity.

Steady-states of Eq.(1) obey Δ = 0, and marginal stability along the manifold is determined by the condition 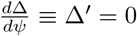 (Fig. S3A). Crucially, the evolution of *ψ* along the manifold is assumed to be dramatically slower than the convergence toward the manifold (see below for cases where this assumption does not hold).

We next apply the simple dynamical rule of Eq.(4) to analyze the network’s response to external stimuli and to study how a continuum of persistent neural states emerges in the trained networks.

#### 2.2.1 Response to external input

Assuming that the learning is successful and a manifold attractor emerges in the recurrent network, the drift along the attractor is negligible (see Section 2.2.2). Upon introduction of external input, the continuum of persistent states collapses to a single fixed-point attractor (Fig. 3, Fig. S3G-H) and the neural state, and hence the representation of the feature *ψ*, begins to evolve according to Eq.(4).

Figure 3A depicts an example in which after training an heterogeneous ring manifold we update the bump position by introducing a weak stimulus in a direction of *ψ*_1_ = *π*/2. The update dynamics are in agreement with Eq.(4) (Fig. 3B, compare colored points with the theoretical prediction in dashed lines). Here, the heterogeneity level *g* only affects dynamics via the effective time constant *τ_eff_*, and not via the tangential error, which is given by Δ = –*ϵA*^-1^ sin(*ψ* – *ψ*_1_) (red curve in Fig. 3B, see also Methods and [Ben-Yishai et al., 1995]). For a ring manifold, we calculated the effective timescale and find that it is given by *τ_eff_* ≈ *τ*(1 – *β*(*g*))^-1^, with *β*(*g*) accounting for the effects of synaptic heterogeneity. This factor is connected to the correlation among the individual neuronal gains:

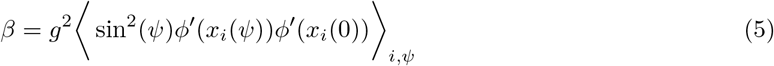

with *ϕ′*(*x_i_*(*ψ*)) being the gain of the neuron *i* at location *ψ* on the manifold and with 〈.〉_*iψ*_ denoting average over all locations and all neurons (Methods). In networks lacking heterogeneity, the factor *β* is zero and it increases monotonically with the heterogeneity level. We therefore find that the larger the amount of synaptic heterogeneity in the trained network, the slower the response to the external input is (Fig. 3C-D).

We next sought to investigate how the dynamics along the manifold are affected by its geometry. For an arbitrary manifold, in contrast to a ring manifold, the time required to update the internal representation from one location to another depends on the initial and final locations and not solely on the difference between them (Fig. 3E-G). This dependency on the specific location on the manifold can be decomposed to the dependency on the geometry of the manifold and the input, as captured by the tangential error, and on the effective timescale which varies along the manifold (Fig. 3F). Figure 3H shows examples of the calculated timescales along various shape of manifolds (see derivation in Methods). Interestingly, we find that the effective timescale tends to be slower at locations along the manifold that are represented by a higher total firing rate (peaks at Fig. 3H; Fig. 2A). This result provides a quantitative support to the intuition that the more tuned the internal representation is (i.e. the larger the amplitude of the bump), the harder it is to update its location and, therefore, the more slowly it responds to external input.

#### 2.2.2 Build-up of a continuum of persistent neural states

We next ask how a continuous internal representation emerges from sampling discrete points on the manifold. In our model, we train the network by sampling *M* points of the manifold *ψ*_1_…*ψ_M_*, which enforces these points to become fixed points of the neural dynamics (Fig. 4A, Methods). However, the neural states at unforeseen values of *ψ* are not persistent and tend to converge to one of the learned fixed points (Fig. 4B,C). While until now we assumed that the number of sampled points is large (1 ≪ *M* ≪ *N*), in this section we consider finite number of sampled points. Specifically, we are interested in assessing how quickly the attractiveness of individual points diminishes with more samples added (Fig. 4B,C), prompting emergence of a continuous manifold, and how the heterogeneity level modulates this effect (Fig. 4B-C). According to Eq.(4), this is equivalent to analyzing how quickly the slope of the tangential error at the sampled points approaches zero. We find that interpolation toward a continuum of the unforeseen values of the feature *ψ* happens very quickly with the number of samples.

We quantify this by analyzing the trained ring manifold (Fig. 1B), which is especially amenable to a full analytical treatment. Here, we show analytically that the rate at which the on-manifold dynamics approaches marginal stability, and hence a continuous representation of the feature, depends on the decay rate of the principal components of the neuronal representation. Specifically, linearizing Eq.(4) around the sampled points yields the eigenvalue of the linearized dynamics:

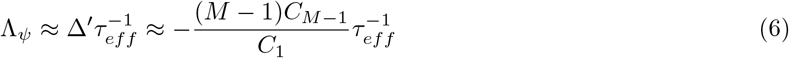

where *C_k_* denotes the *k^th^* score of the neural activity correlation matrix, or, equivalently, variance explained (VE) by the *k^th^* PCs of the neuronal representation (see Eq.(77) in Methods for the exact equation, including small M). We next calculated the decay in VE with the PC number. Figure 4D shows that the VE decays fast with successive PCs, even for high synaptic heterogeneity. Therefore, as the factor 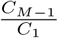 decays fast with *M*, exponentially in case of smooth correlation functions ( e.g. [Katznelson, 2004]), the derivative of the tangential error gets smaller, and the dynamics along the manifold becomes marginally stable, with no attractive or repelling forces present (Λ_*ψ*_ ≈ 0, Fig. 4B-C,E-F). The analytical derivation is in good agreement with the simulations (compare solid lines with circles in Fig 4E-F).

Finally, another indication for convergence to a continuous attractor is a small drift velocity between the sampled points (see Eq.(4)). This is verified numerically in Fig. S5A, showing that the drift velocity is small and also decays exponentially with the number of points.

While it is not straightforward to generalize Eq.(6) to a manifold with arbitrary geometry, such as those presented in Fig. 3, we find that also in such cases the continuity along the manifold is obtained very quickly, exponential with the number of sampled points (Fig. S5B-C). We thus conclude that the rate of approaching continuity of feature representation in the trained networks is extremely fast, even with heterogeneity in synaptic connectivity and asymmetries in the neural representation.

### 2.3 Convergence toward the manifold attractor

Neural dynamics must be robust to perturbations and stimuli that push the neuronal activity away from the manifold attractor [Durstewitz et al., 2000]. Here, we distinguish between the neuronal activity that affects the bump’s dynamics, i.e., its location and amplitude, and the vast majority of activity, which is orthogonal to decoder plane and, as long as it is stable, does not affect the bump’s dynamics (Fig5A).

**Figure 5:**
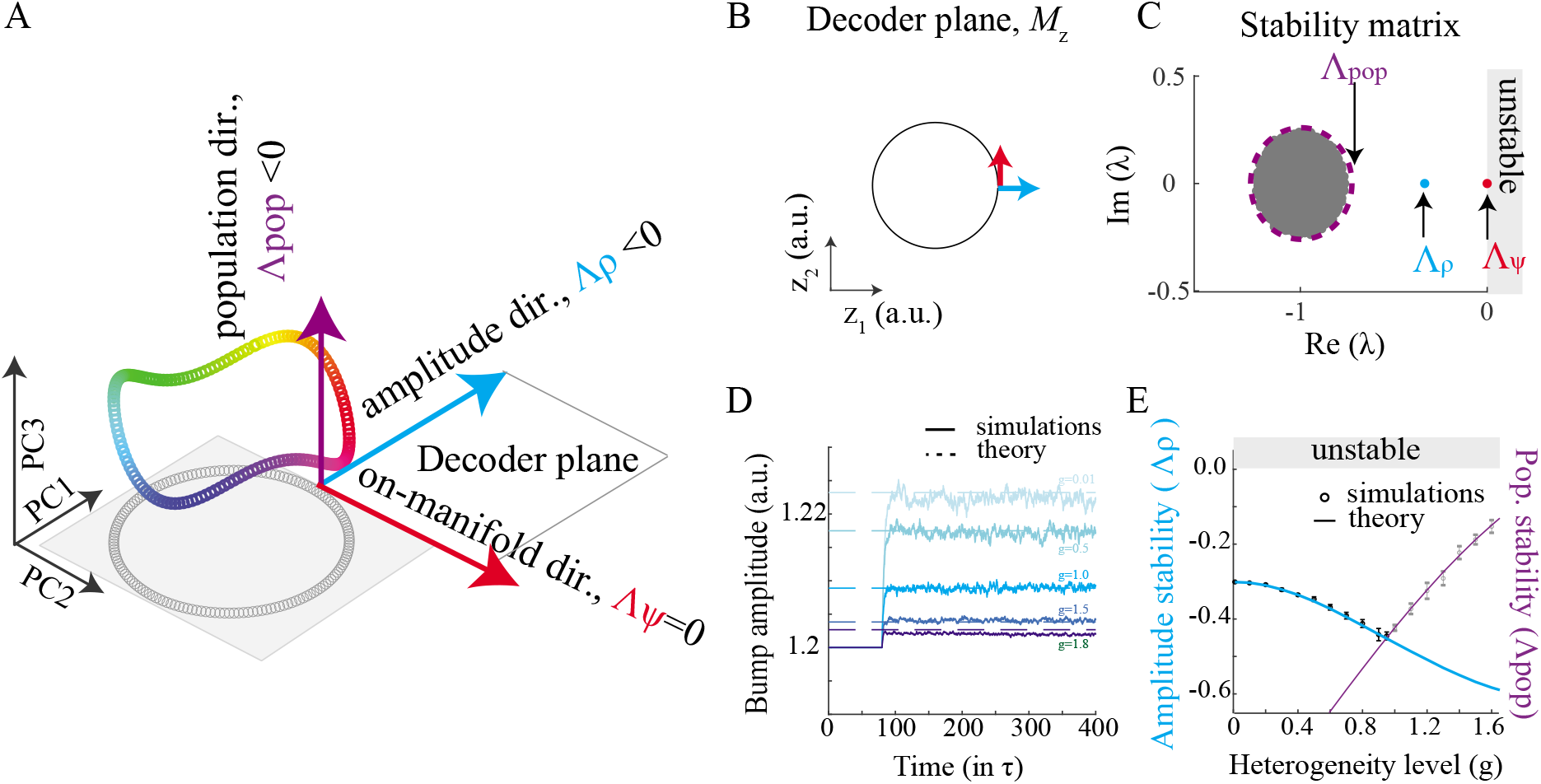
The effect of synaptic heterogeneity on the convergence toward the manifold. **A-C.** Illustration of the space of neuronal activity and of subspaces used to determine the stability of the manifold and their expected properties. **A.** Linearizing the dynamics around the manifold results in modes that affect the stability of the on-manifold (red) and amplitude (blue) directions, as well as other modes of the neural dynamics (purple) that are invisible to the decoder plane and, hence, do not affect the bump’s dynamics. For a ring manifold the decoder plane is aligned with the first 2 PCs. **B.** Projection of the manifold on the decoder plane and illustration of the amplitude and manifold directions for which we linearizate. We expect a zero maximal eigenvalue in the manifold direction (Λ_*ψ*_ ≈ 0 for marginal stability, Eq.(6)), and negative maximal eigenvalues in the amplitude (Λ_*ρ*_) and the other population directions (Λ_*pop*_). **C.** Spectrum of the linearized dynamics of Eq.(1) (imaginary and real parts of the eigenvalues, λ’s, of the stability matrix in Eq.(14)) around the trained points. We denote the maximal eigenvalue at each direction by Λ. Dynamics are unstable if one or more eigenvalues cross zero (gray area). **D,E.** Synaptic heterogeneity stabilizes the amplitude direction. **D.** Bump amplitude against time following a perturbation at time 80*τ* for different levels of synaptic heterogeneity. The larger *g* is, the more stable is the amplitude and fluctuations are damped. **E.** Second largest eigenvalue of the stability matrix (obtained from simulations) against heterogeneity. Mean± sd of simulations of 10 network realizations. Blue: predicted (largest) maximal eigenvalue in the amplitude direction ( Λ_*ρ*_, see Methods 6.7.3). Purple: predicted maximal eigenvalue of the population direction; see Eq.(15) for Λ_*pop*_.

For a heterogeneous ring manifold, we find that synaptic heterogeneity stabilizes perturbations in the bump’s amplitude (Fig. 5D,E). Here, a persistent state on the manifold is perturbed by a noisy external input in a direction that affects the bump’s amplitude. In good agreement with mean-field theory (Methods), both static perturbations and fast noisy fluctuations are damped more when synaptic heterogeneity is increased. This effect is determined by the second largest eigenvalue of the linearized dynamics of Eq.(1) (Fig. 5C), corresponding to the amplitude direction for which simulations and theory are compared in Figure 5E (blue line, see Methods). Qualitatively, this additional stabilization could be attributed to an increase of neuronal activity, causing more neurons to saturate and become less prone to perturbations.

The situation is more complex for a general geometry, where we demonstrate that the convergence toward the manifold might be compromised and even ruined. Figure 6 depicts an example of a family of manifold geometries, parametrized by a continuous parameter *h* so that *h* = 0 corresponds to a perfect ring and positive or negative h implies gradual deformation. Here we focus the analysis on points with reflection symmetry (Fig. 6A), as we find that instability tends to show up at these points (Fig. S6, Methods).

**Figure 6:**
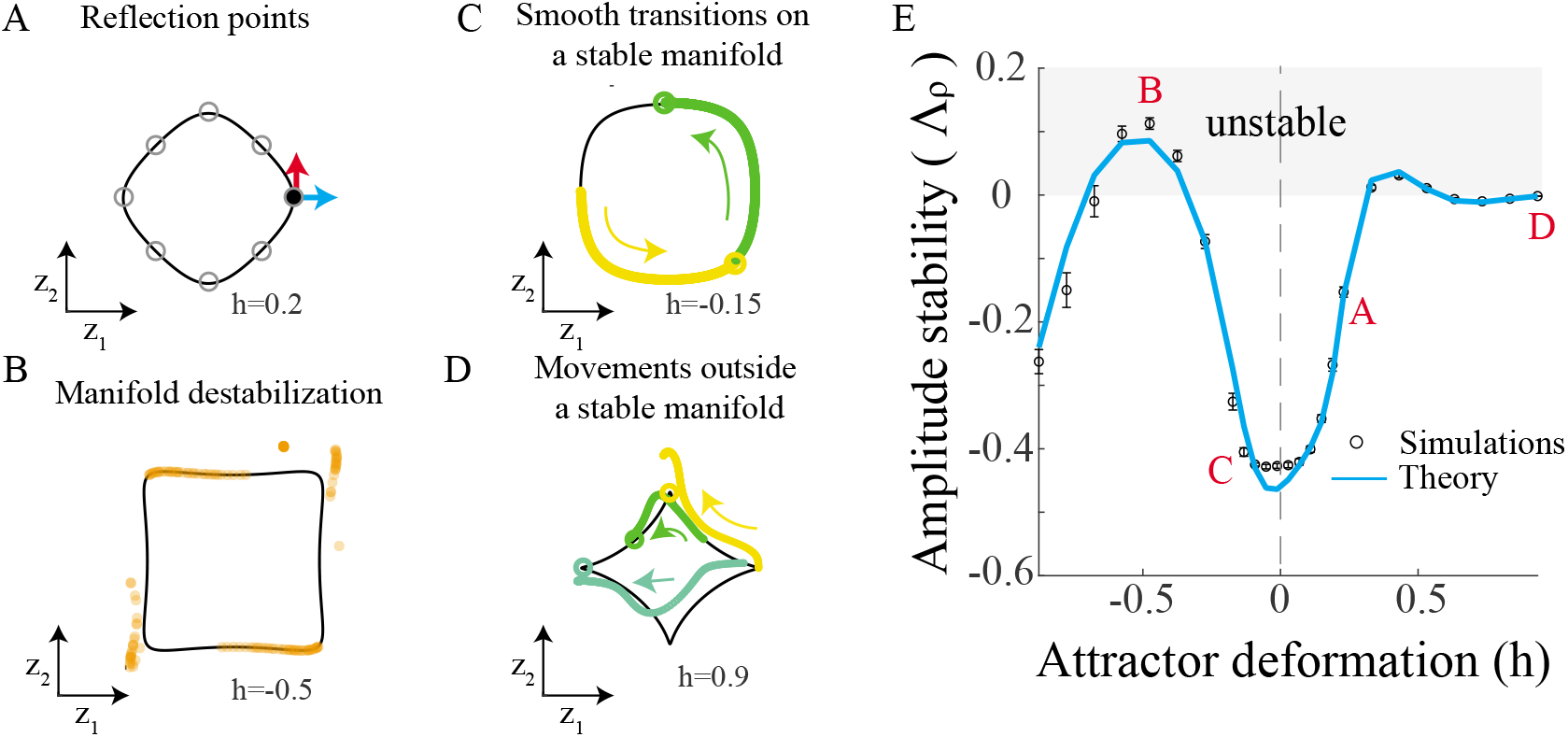
Geometry of manifold attractors controls the convergence in the amplitude direction. A family of trained manifolds, parameterized in Eq.(7), with *h* = 0 corresponds to a ring manifold. **A.** Projection of the manifold on the decoder plane and illustration of the amplitude (blue) and manifold (red) directions at 0 radians (black point) for which we linearize. In general, we analyzed stability at reflection points (gray circles). Black curve: predefined trained manifold **B.** Destruction of a manifold attractor due to instability of the amplitude direction. A snapshot of the neural activity, where all points initialized on the trained manifold, projected on the decoder plane, ***z***. In steady state, all points converge to one of four isolated attractors. **C.** Example of a manifold for which the amplitude direction is stable and the internal state is updated smoothly along the manifold (same figure as in Fig. 3E, bottom). **D.** Same as (C), but for a manifold for which the amplitude direction is close to instability (see (E)). In the presence of input the trajectories deviate from the manifold. See also Fig. S4. **E.** Amplitude stability, as measured by the maximal eigenvalue of the stability matrix along the amplitude direction, vs. deformation of the manifold, h. Circles: maximal eigenvalue in the amplitude direction (blue arrow in (A)) calculated from the spectrum of the full stability matrix, Eq.(14) (see Methods). Blue line: theoretical prediction through the low-dimensional mean-field stability (see also Fig. 7). Note that some manifold geometries are unstable, as shown in (B) for h=-0.5.

Figure 6B shows an example of geometry for which the manifold is unstable. Destabilization of the manifold, which is due to amplitude direction (Fig. 6D), can be non-monotonic in both the complexity of the manifold’s shape (in h, Fig. 6D) and in the amount of synaptic heterogeneity (not shown).

If the bump amplitude is close to instability, as in the example depicted in Fig. 6D, the separation of timescales, which Eq.(4) is built upon, no longer holds. Contrarily to the attractor networks in Fig. 3, here a weaker attraction, or equivalently higher susceptibility to input in the amplitude direction, enables input-driven trajectories to leave the manifold. Thus, instead of moving smoothly along the manifold (as in Fig. 3 and Fig. 6C), the neural state jumps toward a new location that is dictated by the external cue (Fig. S4).

The dynamics of the bump, and in particular its destabilization, can be assessed self-consistently using a small number *K* ≪ *M* ≪ *N* of dimensions of neural state (see Methods, Fig. 7). Specifically, these dimensions are the K leading principal components of the neural representation along the manifold (Fig. 7A). Figure 7B-D depicts the number of PCs needed to explain the manifold stability for the example shown in figure 6, together with another example of a family of ellipse manifolds. For a ring geometry (*h* = 0), only K=2 dimensions are needed to predict bump dynamics, one dimension for bump’s amplitude and one for its location (Figs.7Bi,C-D and Fig. S7A). It is only when considering more involved geometry of the manifold that the number of required PCs is larger than two (e.g. Figs.7Bii-iv and 7C-D). However, as long as the deformation of the manifold is small, the dynamics change continuously and moderately with h, and only a few PCs are needed to predict the bump’s dynamics. Remarkably, the embedded dimension of the manifold, measured for example by participation ratio ([Gao et al., 2017], Methods), is not correlated with *K* (e.g. Fig. 7B), suggesting that it is the geometry of the manifold that controls the bump’s dynamics and not the number of PCs needed to reconstruct the manifold. Interestingly, destabilization was typically associated with large K needed to capture dynamics (Fig. 7D), suggesting that destabilization mechanisms and complexity of geometry, as indicated by large K, are interrelated.

**Figure 7:**
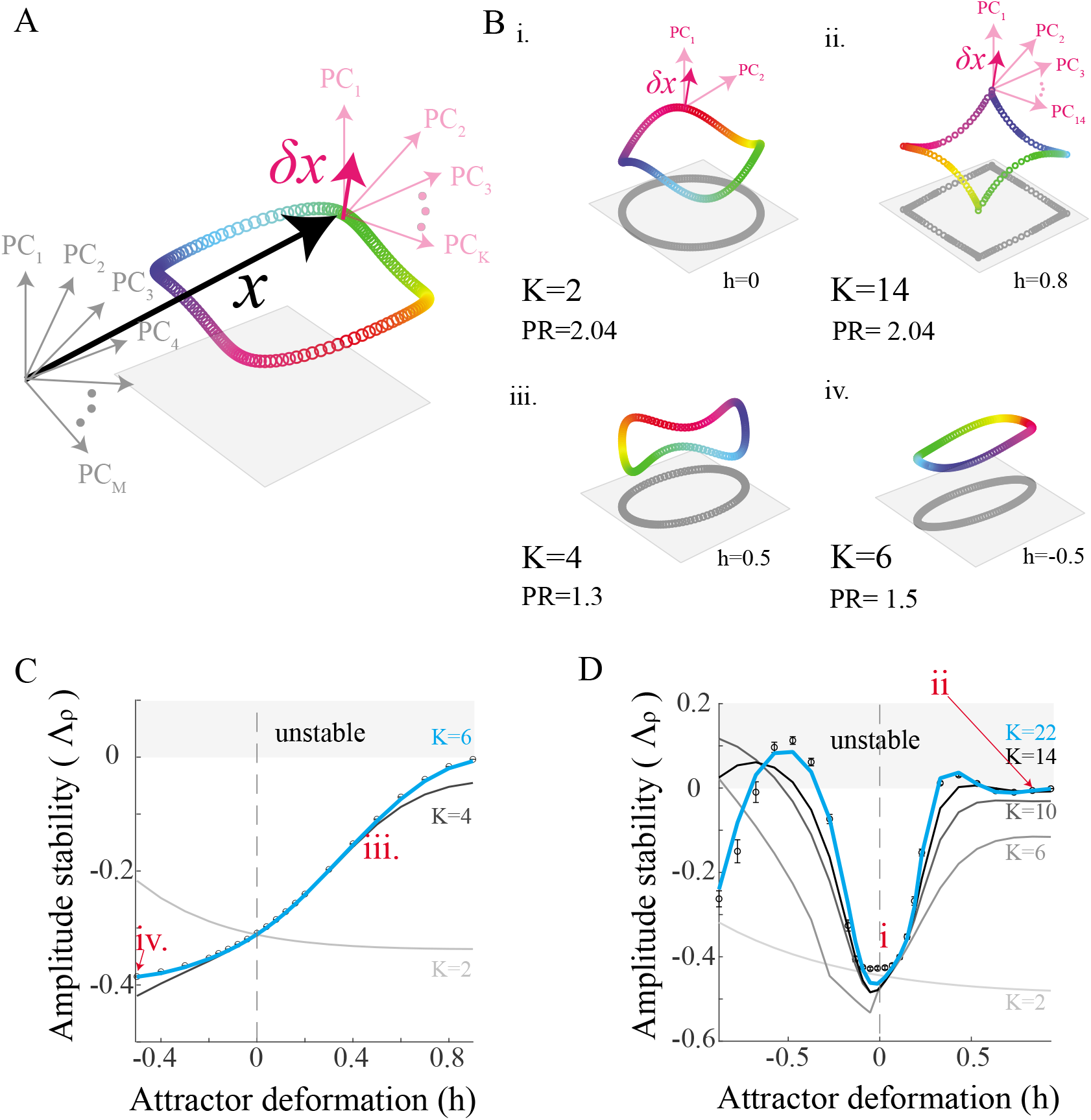
Geometry of manifold attractors determines the complexity of local dynamics in the vicinity of the manifold. **A.** Illustration of the dimensionality reduction in local dynamics: *x* is a point in the N-dimensional space of neural rates. *δ**x*** is a perturbation to ***x*** for which one needs to assess stability. If ***x*** (or equivalently *ϕ*(***x***)) belongs to the training set it can be spanned by the M principal components of this set (grey). Perturbation *δ**x*** can be accurately evaluated in a smaller subspace of *K* out of M principal components (pink). **B.** The number of dimensions *K* needed to accurately evaluate stability is shown along with participation ratio (PR, a measure that estimates the linear dimensionality of the manifold, see Methods) for few representative cases: (i) dynamics in the vicinity of a manifold with ring geometry is of *K* = 2 dimensions, (ii) a manifold with same participation ratio as in (i) but with extremely intricate geometry and thus a high *K*, (iii,iv) examples where participation ratio is smaller than in the case (i), but *K* is higher. **C.** Ellipse manifolds, parameterized as in Eq.(7), with *h* = 0 corresponds to a ring manifold. Amplitude stability is analyzed at 0 radians, as in Fig. 6E. Solid lines: theoretical prediction through the low-dimensional mean-field stability matrix (Eq.(94)) with a cutoff at K=2,4 and 6 PCs. **D**. Same as (C), but for the manifold in Fig. 6. and with higher values for cutoff. Points that were presented in panel (B) are marked in red.

To conclude, we find that it is possible to learn manifold attractors with neither symmetry in synaptic connectivity, nor symmetry in the shape of the manifold. However, the convergent dynamics toward the manifold, and transitions along the manifold, depend both on the geometry of the manifold and the synaptic heterogeneity. For some cases of complex geometry, this convergence is either partially compromised or ruined, resulting in jumps instead of smooth transitions along the manifold, or even in its complete destabilization. The dimensionality of the dynamics in the vicinity of the attractor is determined by the complexity of the attractor geometry, with the special case of ring geometry being especially amenable to analytical formulation and analysis, but otherwise not exceptional.

## 3 Discussion

The hypothesis of computations by manifold attractors relies on idealized symmetry assumption in synaptic connectivity [Fisher et al., 2013], which constitute the theoretical backbone for the emergence of such attractors in recurrent neural networks [Chaudhuri and Fiete, 2016]. However, this assumption is inconsistent with heterogeneous synaptic interactions and single-neuron properties, as well as with the diverse neural responses that are widely observed in various brain areas. Furthermore, as a result of this symmetry in the connectivity, such symmetric-connectome networks can support only a limited repertoire of internal representations, which must themselves have symmetric geometry. Thus, the validity of the symmetry assumption in real biological systems is highly speculative.

We argue that symmetry is not required to produce an approximately continuous manifold in recurrent networks. Instead, training networks results in manifolds of states that can be considered persistent for any practical purpose, without symmetry in the recurrent interactions or in the neuronal representation. Specifically, exponential decay of attracting forces along the manifold is predicted analytically for tractable cases and verified numerically to hold for more involved settings. A relatively small number, order of ten, of learning samples, nulls any motion on the manifold.

Besides the ability to maintain persistent states, functional manifold attractor must be responsive to external input in a tractable way. We found that the response to external input is determined primarily by attractor geometry, and more specifically by its projection into decoder plane. Namely, to make a coarse prediction of how the memorized feature will evolve upon introduction of input, one does not need to know about the internal connectivity of the neural network. For more accurate estimation of the input-driven dynamics, another factor is needed, which is an effective timescale that modulates the motion along the manifold and is expected to affect the ability of the network to perform angular integration (see below). This factor does depend on internal connectivity and can be devised self-consistently in our model, leading to a fine and accurate prediction of dynamics (see Fig. 3).

Finally, in contrast to the bump’s location, which must be responsive to stimuli, its amplitude is expected to remain approximately indifferent to the external input and perturbations [Durstewitz et al., 2000]. In line with this requirement, we found that amplitude stability is preserved even when symmetry of neuronal representation is compromised (Fig. 6). However, we do find that destabilization of the bump’s amplitude is a limiting factor in the emergence of manifold attractors. When geometry of neuronal representation becomes overly complex, the bump’s amplitude may become unstable. Such a complexity is associated with increasing number of dimensions of the neural representations that is required to predict dynamics in the vicinity of the manifold (Fig. 7). This result suggests that the geometry of putative manifolds in the brain can be more involved than a ring, but not overly involved to maintain a stable representation.

### 3.1 Heterogeneity in networks that memorize and path integrate

Storing spatial information and path integration have been hypothesized to rely on the concept of computations by manifold attractors. Neurons in brain areas that support these computations are known to be highly heterogeneous [Romo et al., 1999, Finkelstein et al., 2015, Murray et al., 2017, Fisher et al., 2019, Chaudhuri et al., 2019]. Neuronal activity in symmetric-connectome networks, however, are inconsistent with these studies, as neurons in these models do not exhibit any heterogeneity. In fact, adding heterogeneity to these networks results in a systematic drift of the memory and to a failure to integrate and accurately respond to external cues (Fig. S1).

A few studies have explored the ability of manifold attractors to cope with diversity in neuronal tuning and heterogeneity in synaptic connectivity. It is possible to reduce the drift of the internal representation by adding a slow component to the dynamics, such as short-term facilitation [Itskov et al., 2011, Hansel and Mato, 2013]. In trained networks, such dynamical mechanisms may account for rapid alterations to connectivity for which training proves too slow. This possibility is left for future work. Renart et al.[Renart et al., 2003] showed that homeostatic mechanisms can homogenize the network despite considerable heterogeneity in single-cell and synaptic properties. Our results indicate that even with heterogeneity overcome, its dynamical consequences persist: converging forces toward the manifold in the amplitude direction increases with the amount of heterogeneity (e.g. Fig. 5) and the input response slows down (Fig. 3). Furthermore, the notion of homogenization becomes irrelevant once the requirement for symmetric geometry is relaxed, and neural states become manifestly diverse (Fig. 2, 7).

Instead of relying on rotational symmetry to construct a continuous attractor, a different approach was taken by [Mastrogiuseppe and Ostojic, 2018]. The recurrent connectivity in this model is based on a Hopfield network [Hopfield, 1982], and symmetry in strength of two attractors translates into continuity. However, this approach leads to a limited repertoire of tuning curves, differing only by their amplitude, and does not cope with asymmetry in synaptic connections. Furthermore, adding randomness to the recurrent connectivity in this model shatters the continuous attractor [Mastrogiuseppe and Ostojic, 2018]. This shattering can be mitigated by training, and is tractable by our analysis (see Methods).

Manifold attractors also emerge without training in heterogeneous networks of gated units [Krishnamurthy et al., 2020]. In contrast to the 1-D trained manifolds studied here, the dimension of these manifold is macroscopic. Moreover, it is unclear how to construct the external input to these networks in order to update the representation along the manifold.

In application to systems that track external inputs and integrate angular velocity, such as the head-direction system [Hulse and Jayaraman, 2020], our analysis can be used to obtain the maximal tracking velocity. As the response to external input slows down in the presence of synaptic heterogeneity, we expect the maximal tracking velocity to decrease with synaptic heterogeneity in the network. As heterogeneity is correlated to diversity in tuning curves in our model, our work suggests that integration is impaired in networks where neuronal responses are overly diverse.

In the fly’s head-direction system, the network supporting the integration of idiothetic and allothhetic signals is assumed to include 10-50 neurons [Hulse et al., 2020]. In this and other [Simony et al., 2008] small networks, almost any deviation from a symmetry assumption, such as heterogeneity in tuning curves and in the connectome, is catastrophic for the computational capabilities of the network. While here we applied a recurrent autoencoder paradigm (Fig. S3 and Methods) to analyze manifold attractors in large networks, in which mean-field estimates are obtainable for Eq.(4), the paradigm itself is valid for networks as small as a few neurons.

Finally, noise accumulation is a limiting factor in working memory and in integration systems [Burak and Fiete, 2012], and is attributed to the marginal stability of the manifold direction. Our analysis implies that not all the marginally stable manifolds were born equal: we found that both synaptic connectivity and the geometry of neuronal representaton affect the drift along the manifold. It will thus be interesting to explore how these effects generalize to diffusive dynamics.

### 3.2 Beyond symmetrical geometry in manifold attractors

Symmetric-connectome models can support only the representation of manifolds with symmetry in their geometry, in which the tuning curves are identical, the projections on the two leading principal components (PCs) are circular, and population rate is constant along the manifold (1A). However, it is unclear that this is the case for putative manifolds in the brain. For example, in the head direction system [Rubin et al., 2019, Chaudhuri et al., 2019] leading PCs do not seem to feature such a perfect circular shape. Moreover, recent studies suggest that the richness of the environment or locations of learned rewards affect the geometry of the manifold [Boccara et al., 2019, Low et al., 2020].

Our work provides a link between the geometry of the manifold and its dynamical properties. While estimating the geometry of the manifold can be challenging, for example due to small number of recorded neurons or sampling biases [Rubin et al., 2019, Chaudhuri et al., 2019], our work suggests that estimating the manifold’s geometry, and not only its topology, is essential for predicting the computational properties of the network. In particular, our work suggests that the dynamics in the vicinity of locations that are represented by a higher total firing rates (Fig. 2) are less responsive to updates (Fig. 3) and more prone to perturbations (Fig. S 6), with potential implications for computations such as tracking and integration.

Finally, recent computational works show that a manifold attractor emerges in networks with heterogeneous connectivity when recurrent network is trained to integrate angular velocity [Cueva et al., 2019, Sorscher et al., 2020]. Interestingly, the authors in [Sorscher et al., 2020] found a distorted 2D manifold structure when they trained networks to path integrate. Our work provides a theoretical understanding of the connection between the neuronal representation, such as diversity in neuronal responses and the geometry of the manifold, and the emergence of a marginal direction and the dynamics in the vicinity of the attractor in such trained networks.

### 3.3 Analytical theory of trained neural networks

While continuous attactors were observed numerically [Seung, 1998, Seung et al., 2000, Mante et al., 2013, Sorscher et al., 2020, Cueva et al., 2019, Maheswaranathan et al., 2019], it is not clear how they emerge from a finite number of training examples. Here we established a link between interpolation capabilities and the spectrum of neuronal activity. Specifically, Eq.(6) connects the decay rate of the PCs of neuronal tuning curves to the rate of approaching a marginal stability. From a signal processing perspective (e.g. [Shannon, 1949]), this result can be interpreted as quantifying frequency *aliasing*. The decoder samples the leading Fourier mode (i.e sin *ψ*, cos *ψ*). If a finite number of samples, *M*, used for learning, then the *M* – 1-th spectral mode is not orthogonal to the leading mode and it folds on the desired decoder (Eq. (6)). Interestingly, a recent study made a connection between how fast different modes are learned and the complexity of the neural code [Bordelon et al., 2020]. Similar methods might apply to our framework and, to this extent, generalizing our theoretical result on the fast convergence rate toward marginal stability along the manifold (Eq. (6)) to more general topology could prove insightful. Furthermore, extrapolation, which is known to be remarkably harder than interpolation, can be also analyzed from this viewpoint: when some restricted interval [*ψ*_1_, *ψ*_2_] is used for learning instead of the entire domain [0, 2*π*), the desired spatial Fourier mode loses its orthogonality to virtually all other modes and not just to the *M* – 1-st one, resulting in poor extrapolation.

Training large neural networks to have a handful of discrete fixed-point attractors results in low-dimensional dynamics [Rivkind and Barak, 2017]. Yet, it was not clear how this result generalizes to continuous attractors, and specifically whether the dimension of dynamics becomes infinitely large at the limit of a large number of training points. Here, we found that the dynamics in the vicinity of continuous neural manifolds is approximable by a small number of dynamical modes, much smaller than the number of training points, and is related to the leading principal components of static neural representation along the manifold. Interestingly, we observed numerically that destabilization of manifold attractor is attributed to growing number of PCs needed to explain the dynamics. Future work may focus on solidifying this relation analytically.

Finally, synaptic heterogeneity has a regularizing role in the trained networks. Indeed, increasing heterogeneity stabilizes the solution in our model (Fig. 5), in a way that is reminiscent of the ridge parameter in ridge regression. Moreover, training with synaptic heterogeneity leads to an increase in contraction toward the sampled points on the manifold (Eq.(6)), and of contraction in the direction orthogonal to the manifold (Fig. (5)). This contractive flavor of synaptic heterogeneity in the network is reminiscent of the role of noise in denoising and contractive autoencoders, methods that are used to capture local manifold structure of data [Alain and Bengio, 2014].

In summary, our work shows that continuous attractors can cope with a large amount of synaptic heterogeneity and asymmetries in the geometry of the attractors, allowing to construct mechanistic models of manifold attractors in the brain and predict the dynamics from their internal representation.

## 4 Acknowledgements

We would like to thank Larry Abbott, Ehud Ahissar, Arseny Finkelstein, James Fitzgerald, David Hansel, Ann Hermundstad, Brett Mensh, Sandro Romani and Inbar Saraf-Sinik for their valuable feedback. A.R. is hosted by Ehud Ahissar for postdoctoral training in Weizmann Institute of Science.

---

**Figure S 1:**
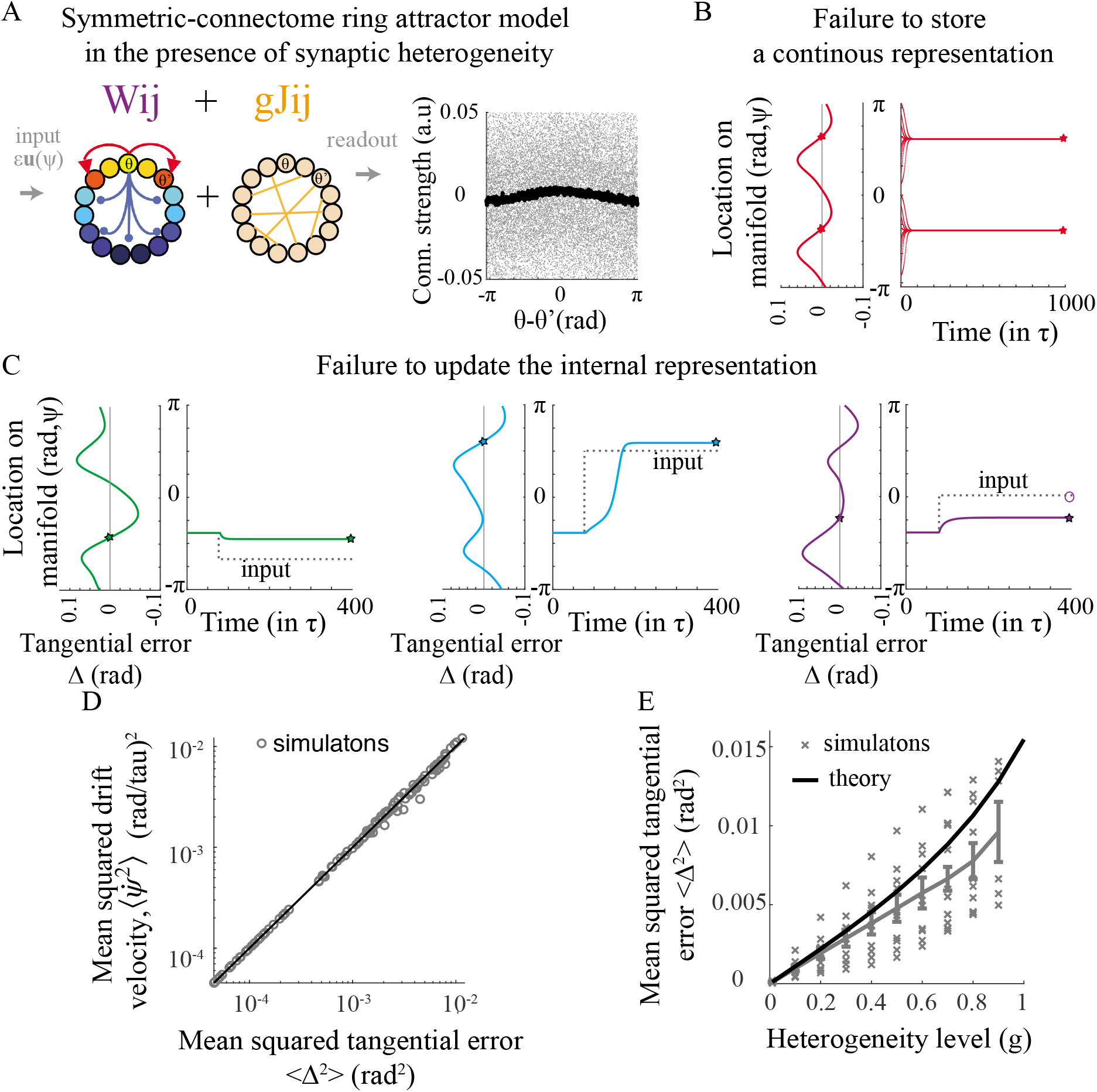
Failure modes of symmetric-connectome ring attractor network in the presence of synaptic heterogeneity. **A.** Right: Cartoon of symmetric-connectome ring model in the presence of synaptic heterogeneity (g=1). The structured part is as in 1Ai and is unlearned. Left: Connectivity strength vs. distance in neuronal preferred directions. **B.** Tangential error vs bump’s location (left) and the bumps location vs. time (right) for the network in (A). Due to the heterogeneity the bump drifts toward one of the two stable fixed points in a few time steps. *N* = 1000, *g* = 1,e =0. **C.** Response of the network in (A) to external input. Same as (B), but with *ϵ* = 0.04. **D.** Mean squared drift velocity, averaged over initial conditions of 160 uniformly sampled points from the manifold, vs. Mean squared tangential error (see Eq.(4)). Each of 100 points corresponds to a combination of hyperparameters *g, A* times five random seeds (Methods). **E.** Mean squared tangential error vs. heterogeneity level. Theory: Eq. (107), developed for small *g*, fits well the simulations for small g and start to deviate from the simulations for *g* ≈ 0.4; *N* = 4000.

**Figure S 2:**
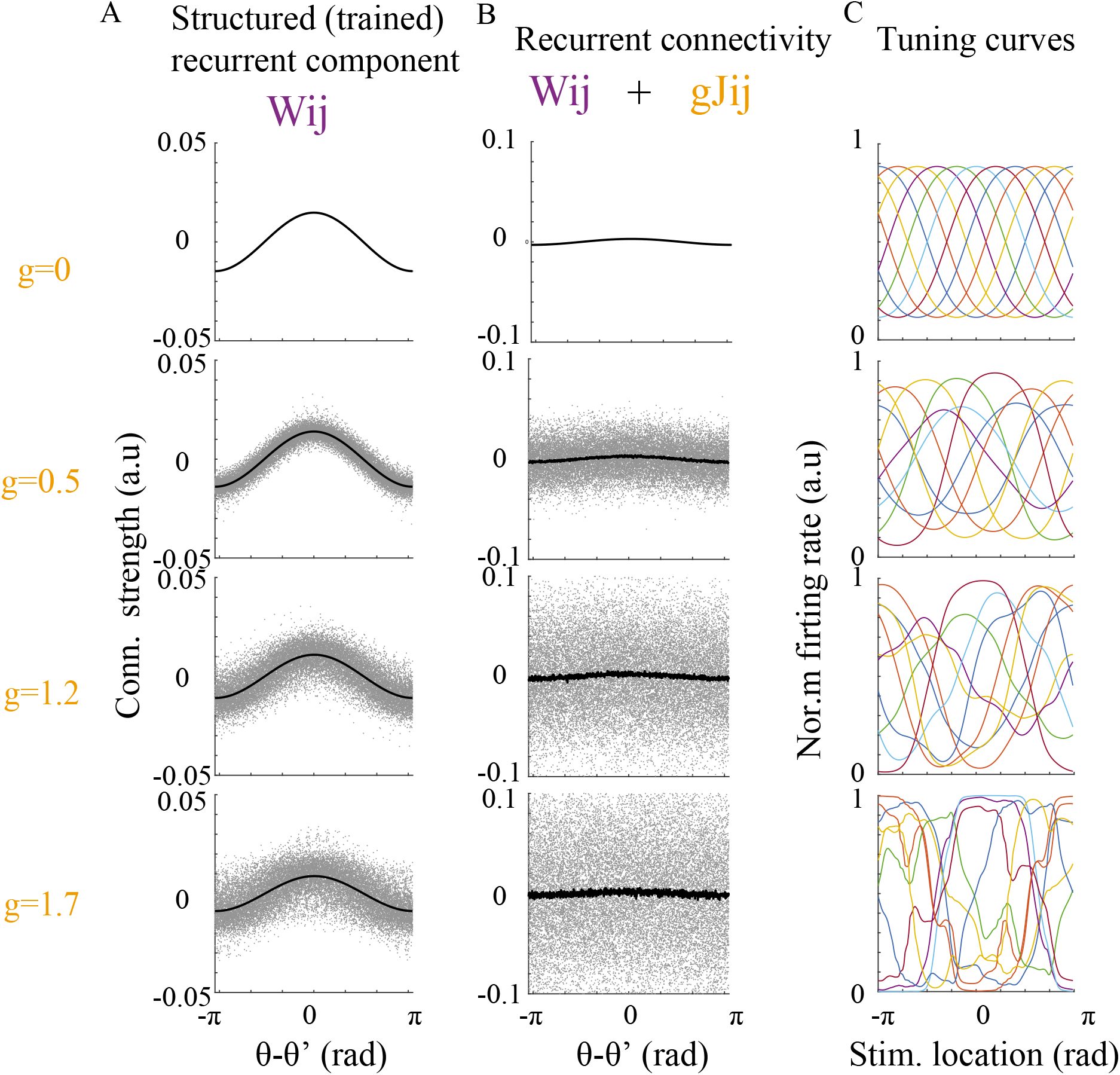
Recurrent connectivity and tuning curves in trained ring attractor networks. **A.** The structured component of the trained network. **B.** The full recurrent connectivity (note the scale difference with (A). Black: average over all neurons with a bin (*θ, θ* + *δθ*) **C.** Tuning curves vs. stimulus location.

**Figure S 3:**
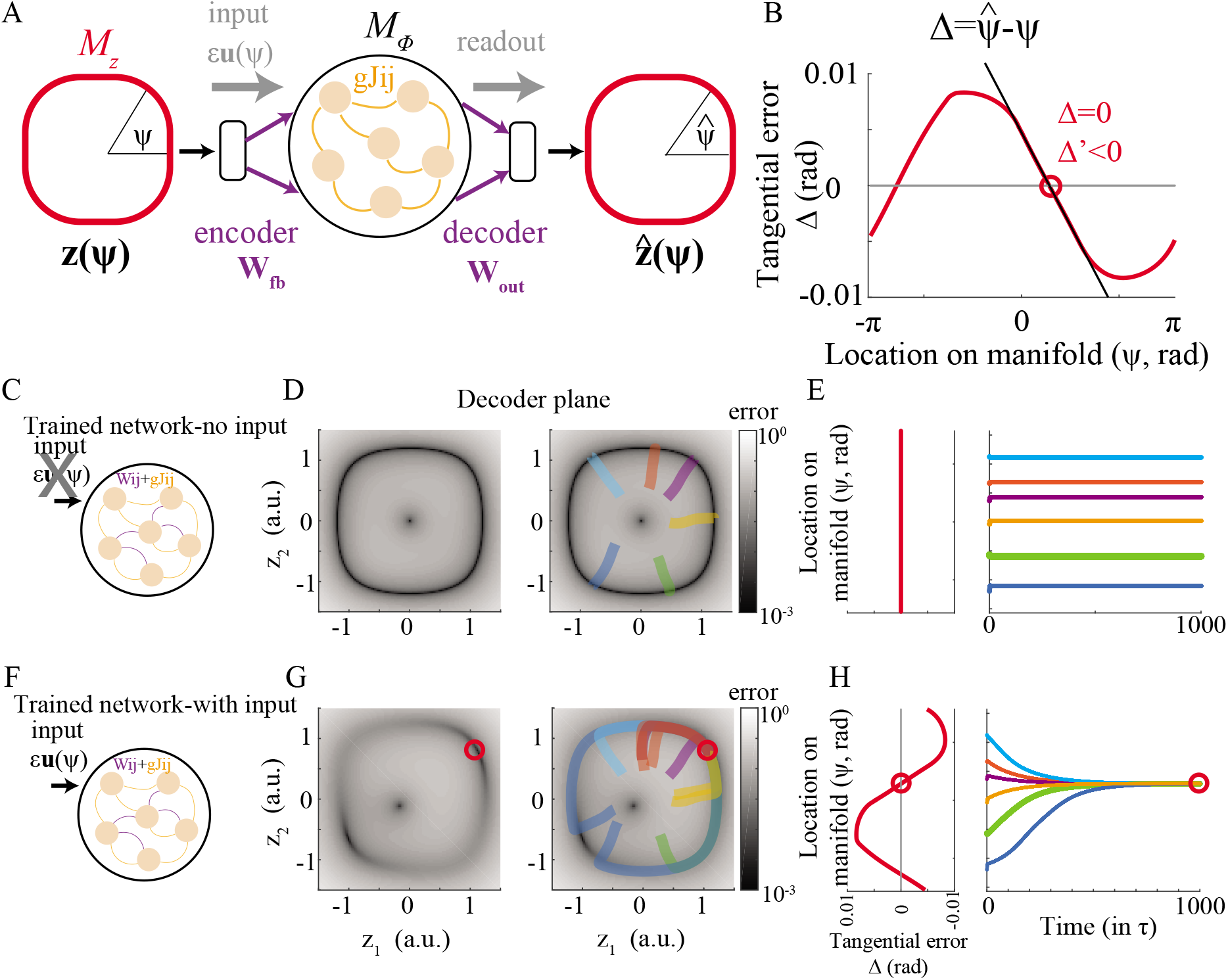
A Recurrent Auto-Encoder framework (RAE) for analysing manifold attractors. **A.** Cartoon of the RAE. The structured recurrent loop (purple lines in Fig1C) is opened and decomposed into an encoder and decoder. RAE is driven by fixed stimulus on the 2D plane ***z*** and the corresponding steady state output *z* is decoded. **B.** The tangential error, between encoded and decoded angles 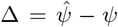. Fixed point of the dynamics are points in which Δ = 0 and stable fixed points (red circles) with Δ′ < 0 (Eq.(4)). **C.-H.** Implications of decoding error on network dynamics. **C.-E.** The trained network in the absence of inputs. **D.** The reconstruction error of the RAE. Left: error is shown in 2D *z* plane. Right: error is superimposed with 6 randomly initiated trajectories of the full system. Dynamics converge rapidly to the closest point on the manifold in which the error is negligible. Note the log scale of the error. **E.** Left: Tangential error Δ vs. location on the manifold. Right: Trajectories in (D) plotted against time. **F.-H.** Same network as in as **C-E**, but with a weak external input (*ϵ* = 0.01) at *π*/4 rad. Here, convergence to the manifold is followed by drift to a single stable fixed point, in which the tangential error vanishes **H**. Decoded angular feature 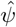 is recovered via 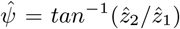. The reconstruction error, 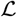 is defined by 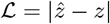

**Figure S 4:**
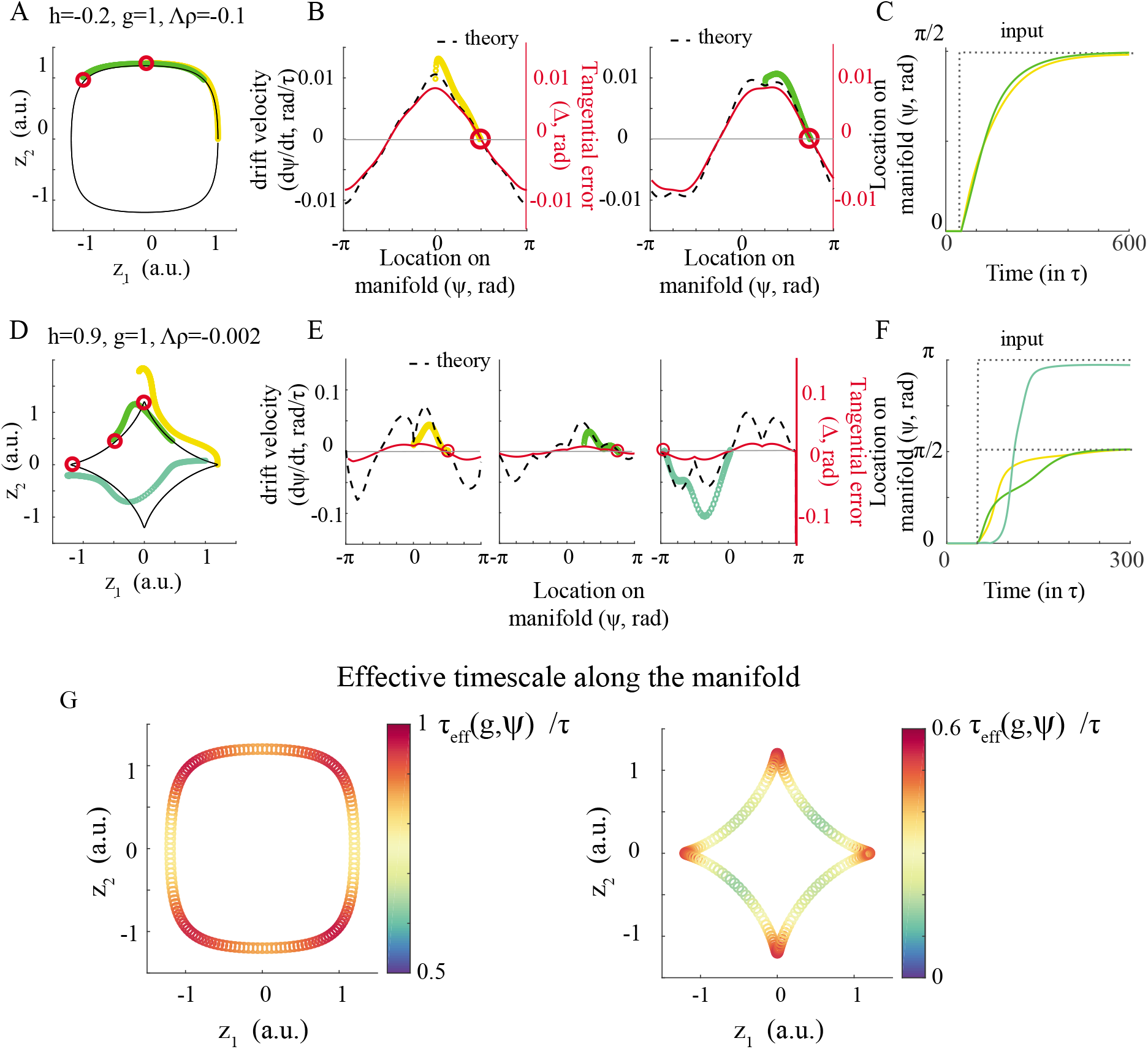
Response to inputs in manifolds for which amplitude direction is not strongly attractive. **A.-C.** Example manifold where amplitude direction approaches instability (maximal eigenvalue along all points on the manifold in amplitude direction is Λ_*ρ*_ = –0.1). Note deviations between theory and simulations due to the lack in timescale separation between the amplitude and manifold directions. **D.-F.** Example manifold where amplitude direction is marginal (maximal eigenvalue along all points on the manifold in amplitude direction is Λ_*ρ*_ = −0.002). Note that transitions are no longer along the manifold. **G.** Effective timescale along the manifold.

**Figure S 5:**
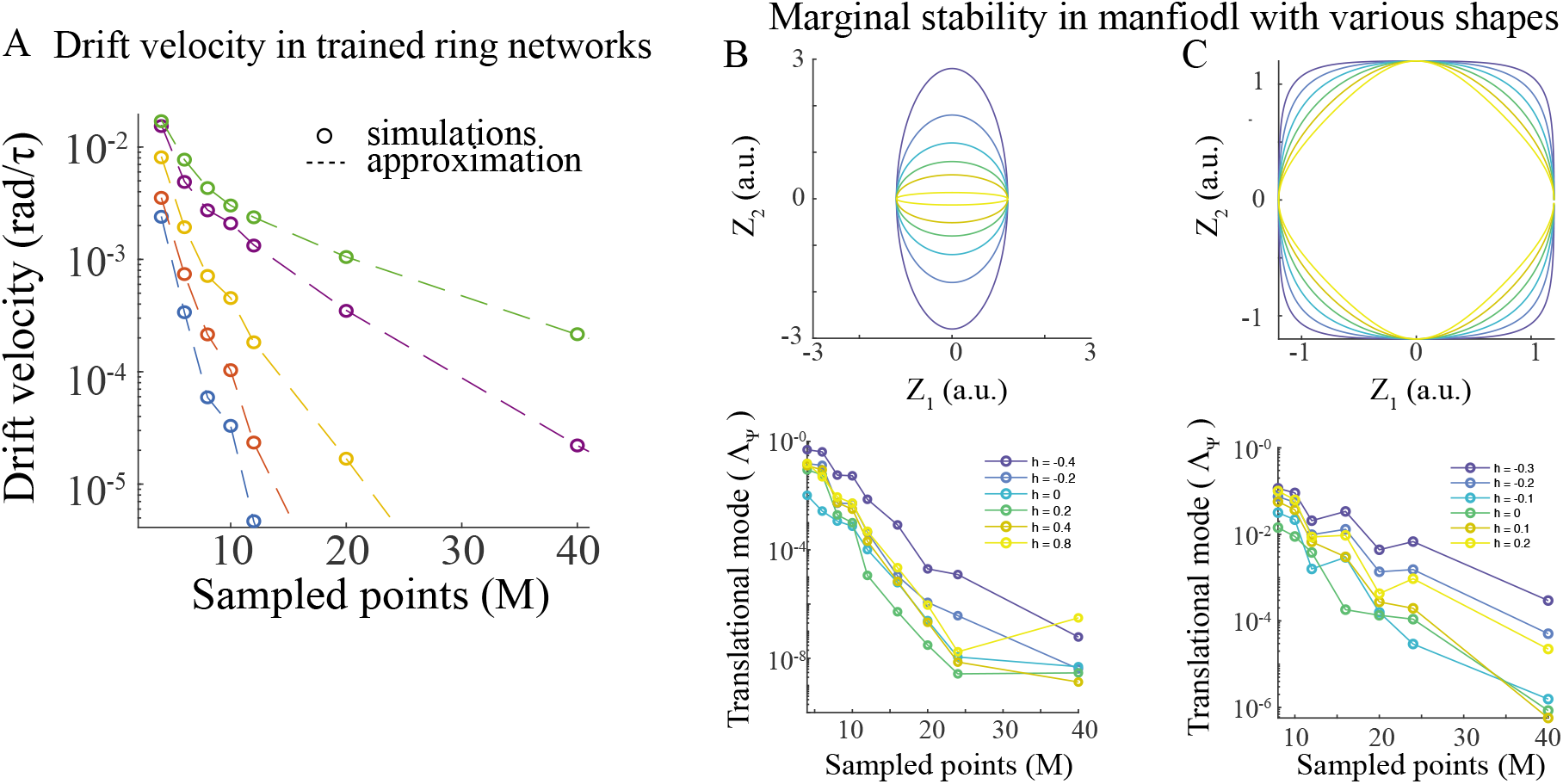
Drift velocity and marginal directions in manifold attractors. **A.** Drift velocity vs. number of sampled points in trained ring manifold. Same data as in Fig. 4F. **B-C.** Build-up of manifold attractor beyond ring geometry. **B** Top: manifold shape projected on the decoder plane. Bottom: Stability along manifold direction for the manifolds in (A). Manifold parameterization: 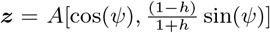 **C.** Same as (B), but for a different manifold, defined by 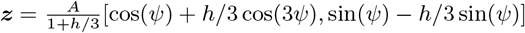.

**Figure S 6:**
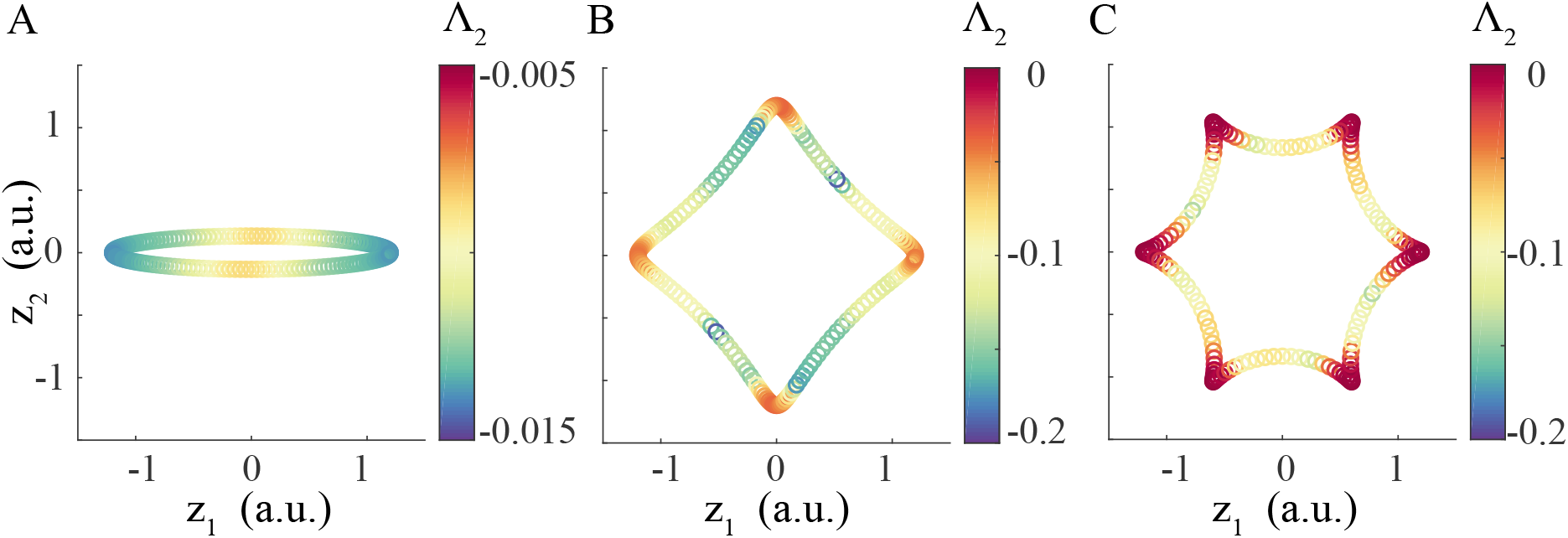
Marginal stability in the amplitude direction tends to occur at reflection points. **A.** The second largest eigenvalue of the stability matrix of the RNN, Eq.(14), plotted along the ellipse manifold. *a* = 2, *h* = 0.8, *g* = 0.6 **B.** Same as (A), but with *a* = 4, *h* = 0.6, *g* = 1. **C.** Same as (A), but with *a* = 6, *h* = 0.9, *g* =1.2.

**Figure S 7:**
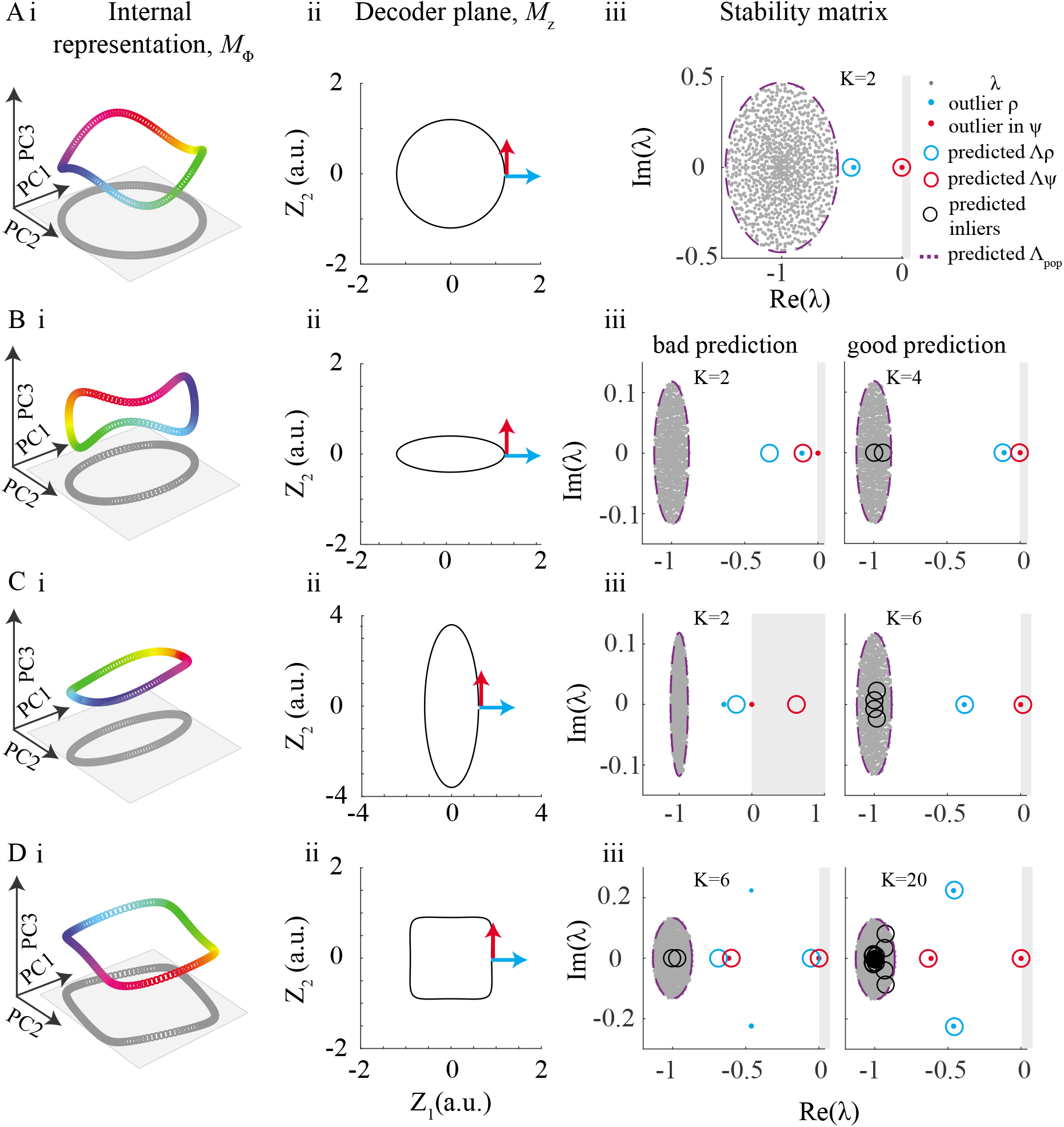
Predicting the spectrum of the stability matrix using the low-dimensional mean-field approach. **Ai.** Projection of a trained ring manifold onto the leading PCs with color coded for the continuous feature. Gray: projection onto the 2 leading PCs. **Aii.** Decoder view of a predefined trained ring manifold. **Aiii.** Spectrum of linearized dynamics of the manifold in (ii). Purple: predicted maximal eigenvalue of the population direction; see Eq.(15) for Λ_*pop*_. Circles: predicted eigenvalues according to the low-dimensional mean-field stability matrix (Eq.(94)), with outlier colored in blue for modes that project to the bump’s amplitude direction and red for the on-manifold direction (see Methods). Here, *K* = 2 principal components suffice to get good mean-field prediction for the outliers. Grey region corresponds to instability (*Re*λ > 0) **B.** Same as (A), but for an ellipse manifold. Here, prediction of the outliers is improved by increasing the number of principal components used for estimating the low-dimensional mean-field stability matrix **C-D.** Same as (B) but for different manifolds

## 5 Methods

### Simulations

All simulations were done using an Euler method with *dt* = 0.1*τ* and *τ* = 1. In some cases we took dt = 1*τ* and checked that similar results hold for smaller *dt*.

### Manifold parameterization

To obtain manifold attractors with non-symmetric geometry, we defined closed curves in the decoder plane. To obtain a family of curves morphing gradually from a ring to a more general geometry we used the following parameterization:

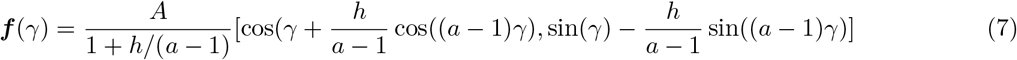

where the normalization 1/*a* + *h* guarantees that the amplitude is *A* at angle *ψ* = 0 radians.

### Training

To train the network we simulated Eq. (16) for 1000*τ* to ensure convergence to steady state and derived the least square solution for ***W**_out_* [Jaeger, 2001]. To simplify the analytical calculations we chose the sigmoidal transfer function *ϕ*(*x*) = *erf*(*x*).

Due to the symmetry of the transfer function, if ***x*** is a steady state of the dynamics in Eq. (11) then −***x*** is a steady state as well. Hence, it is enough to train *M*/2 points at angles 0 ≤ *ψ_m_* < *π* and the other *M*/2 fixed points will emerge at angles *π* ≤ *ψ_m_* < 2*π* due to symmetry. For a manifold attractor with *M* sampled points we thus used 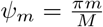 with 0 ≤ *m* < *M*/2.

### Angular feature

*ψ* The angular feature is calculated according to *ψ* = *atan*(*z*_2_/*z*_1_). Note that for a case of general manifold geometry there is no equvalence between angular feature *ψ* and the patameter *γ* that was used to define the curve in Eq. (7).

### External input

In settings with a non-zero external input, the input scaled as ***ũ***(*γ*) = *f*(*γ*)/A to account for changes in the bump amplitude in case of a non-ring geometry. Here, *ũ* is a two dimensional input that projects on the *N* dimensional network, as defined in Section 6.1.

### Recurrent Autoencoder (RAE) Setting

Recurrent autoencoder setting is described in details in Section 6.3 and is depicted in Fig.S 3A. In the RAE setting, the structured part of recurrent connectivity is unfolded and the network is being *driven* by a signal *z*(*ψ*), instead of having this signal regenerated by the recurrent connectivity.

### Tangential error

Fig.S 3A,B depicts how the tangential error, Δ, is obtained. First, a point *z*(*ψ*) from a manifold is encoded as a drive to the dynamics of RAE. Then, the decoded angle is obtained as 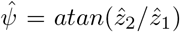. Note that this defines 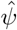 as a function of *ψ*. The tangential error Δ is then given by 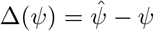.

### Tangential error in the setting with external input

In the case of 0 < *ϵ* ≪ 1, we find that it is sufficient to approximate the tangential error by

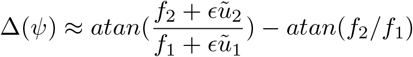

Namely, there is no need to calculate the error using the auxiliary RAE system, and it can be devised solely from ***f*** and the the input ***ũ***. In the specific case of a ring manifold, and for *ϵ* ≪ 1, we get 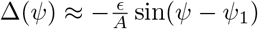.

### Selectivity index and preferred direction

Given the tuning curve of a neuron *k, r*(*θ_k_, ψ*), we calculate the first Fourier component, 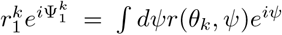, with 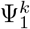 being an estimate for the preferred direction of the neuron and 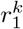 the selectivity index. Selectivity index of zero corresponds to a flat tuning curve and is closely related to a circular variance of one [Cremers and Klugkist, 2018].

### Effective timescale

The effective timescale, *τ_eff_*, is obtained from the gain of RAE. With a detailed derivation of the gain of the RAE given in the next section, and with mean-field estimate for gain obtained using Eq.(81) and Eq.(23), the effective time scale *τ_eff_* follows:

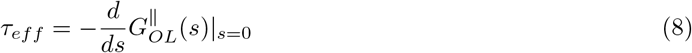

In the specific case of a trained ring manifold, the effective timescale is given by Eq.(70). To compare between theory and simulations in settings of Fig. 3C, we considered the velocity 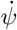 at location *ψ*_1_ = *π*/4 (half-way between the starting and the end point of the trajectory). For each amount of heterogeneity, *g*, the effective timescale was calculated as inverse of ratio between the measured velocity, 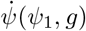, and the velocity in the absence of heterogeneity: 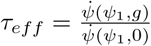.

#### Calculating spectrum of stability matrix

##### Mean-field estimations

In case of a ring geometry, both static correlation function and dynamics in the vicinity of the manifold (in particular the leading eigenvalue of the stability matrix and effective timescale) were obtained self-consistently and compared to simulations. In case of a general geometry, the correlation function was obtained from simulation and dynamics in the vicinity of the manifold was computed self-consistently (Eq.(94)).

##### Dissection between on-manifold and off-manifold directions

For a ring manifold, or at reflection points, we color the outliers of the spectrum of the *N* × *N* stability matrix according to their relation with either the bump’s amplitude (circles in Fig. 6B,C and blue points in Fig. 7A-Diii), or on-manifold direction (red points in Fig. 7A-Diii). Specificaly, we calculated the eigenvalues of 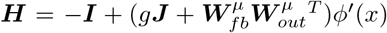, for which *μ* = 1 corresponds to the amplitude of the bump and *μ* = 2 for the on-manifold direction. In the case of the mean-field analysis, we calculated separately the eigenvalues of the normal (amplitude) and the on-manifold direction (see Eq.(94) and the following paragraph).

### Participation ratio

is an estimate for effective dimension of data embedded in high-dimensional space [Gao et al., 2017]. It is calculated as 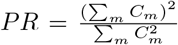, where *C_m_* denotes the *m^th^* score of the neural activity correlation function, or the variance explained by the *m^th^* PC of the neuronal representation.

### Transfer function

To simplify the mean-field calculations, we choose the error function. While this function is very similar to the commonly used tanh, in the case of integration with Gaussian integrals it simplifies calculations and gives an analytical expression for quantities such as the correlation function. Specifically, we use:

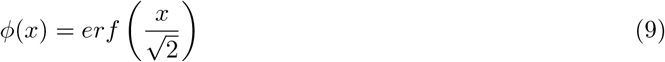

where the error function is defined as 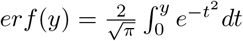. Using the identity 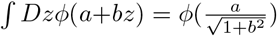 in Eq.(46), we only need to numerically integrate over *θ* and *y*.

### Parameters for figures

Unless written otherwise, the parameters are *A* = 1.2, *g* = 1, *N* = 1000, *M* = 40, *a* = 2, *h* = 0.

- Figure 1: A. *g* = 0, *A* = 1.2. C-D.*g* = 1.2, *A* = 1.2
- Figure 2: A. Black:*h* = 0, *g* = 1. Blue: *a* = 2, *h* = 0.2, *g* = 1. The predefined trained manifold in E-G is given by Eq.(7), with *a* = 2, *h* = 0.2, *g* =1. In A *N* = 16000 and B-I *N* = 4000.

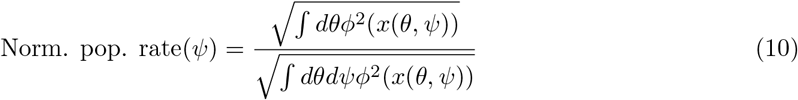
- Figure 3: *ϵ* = 0.01 A-D. *h* = 0. E-G. *a* = 2, *h* = 0.2, *g* = 1. C. N=4000.
- Figure 4: A. *N* = 16000. E-F, *N* = 4000.
- Figure 6: A. N=4000, a=4,g=1, A=1.2.
- Figure 7: B.i.h=0, g=1. ii. a=4, h=0.8, g=1. iii.a=2, h=0.5, g=0.2. iv. a=2, h=-0.5, g=0.2. C. N=4000, a=2, g=0.2, M=40. D. N=4000, a=4, g=1, M=40.
- Figure S1: A-C. Symmetric-connectome ring attractor network (*W_ij_* ∝ cos(*θ_i_* – *θ_j_*), see Methods) in the presence of a random connectivity with *g* = 1. C. *ϵ* = 0.04. D. Hyper-parameters for the scatter plot of drift speed vs. error *A* ∈ {0.5,1.0,1.2,1.5, 2.0}, *g* ∈ {0.01,0.1,0.3,0.5}, times 5 random seeds per setting.
- Figure S2: N=1000.
- Figure S3: *a* = 4, *h* = –0.15, *g* = 1, A = 1.2
- Figure S4: A-C,G left *a* = 4, *h* = –0.2, *g* = 1, *A* = 1.2 D-F,G. right*a* = 4, *h* = 0.9, *g* = 1, *A* = 1.2
- Figure S7: A. h=0, g=0.8. B. a=2, h=0.5, g=0.2. C. a=2, h=-0.5, g=0.2. D. a=4, h=-0.4, A=0.9. g=0.2.

## 6 Details of theory of trained manifold attractors

### 6.1 The network model

We consider a network of N units following the rate dynamics:

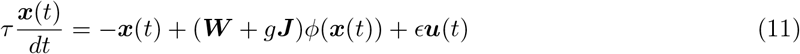

with the neuronal state 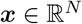 representing the neuronal input and the neuronal rate given by *ϕ*(*x_i_*(*t*)), with a symmetric activation function, *ϕ*(*x*) = –*φ*(–*x*). The recurrent connectivity consists of two components: The random heterogeneous part is represented by i.i.d Gaussian weights 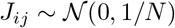 times strength parameter *g* [Sompolinsky et al., 1988]. The other recurrent component is a rank-2 structured part, 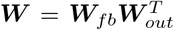, with 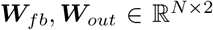. The external input is ***ũ***(*t*) = ***W**_in_**ũ***(*t*), with 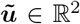, and where we choose ***W**_in_* = ***W**_fb_* for simplicity.

Following the symmetric-connectome ring model of [Ben-Yishai et al., 1995], we choose feedback weights to be 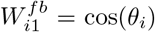 and 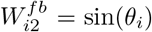, however other choices such as in [Mastrogiuseppe and Ostojic, 2018] give similar results (see also Section 6.9).

The decoder is given by 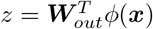, for which the angular feature is calculated according to *ψ* = *atan*(*z*_2_/*z*_1_). The training goal is to obtain ***W**_out_* such that:

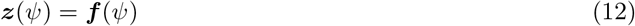

with target 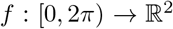 being a curve in 2D. For example, the particular case of a ring manifold is given by ***f*** (*ψ*) = *A*[cos(*ψ*), sin(*ψ*)].

### 6.2 Dynamics in the vicinity of a manifold attractor

For a one-dimensional subset 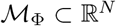 to be a continuous attractor, it is required that in the vicinity of any point 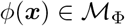, local dynamics will be convergent in *N* – 1 off-manifold dimensions and remain marginally stable in the one remaining direction associated with translations on the manifold (Fig 5A). In the case of the dynamics of Eq.(11), this implies that linearized system:

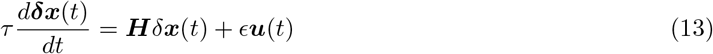

with the stability matrix

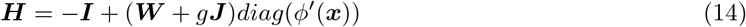

will have a spectrum 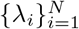 in which λ_1_ = 0 and 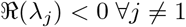. Extending the approach of [Rivkind and Barak, 2017], we argue that training the structured rank-two connectivity matrix affects only a small number of dynamical modes (red and blue circles in Fig. 5C, Fig.S 7), while the rest of the spectrum consists of a bulk of eigenvalues that are associated with random connectivity and are not affected by the structured component. The latter are confined to a circle of radius (purple in Figs. 5C, Fig.S 7, [Ahmadian et al., 2015]):

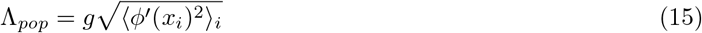

with *x_i_* calculated self-consistently (see below) and 〈.〉_*i*_ average over all neurons. Throughout the work, we assume that the network is in a non-chaotic regime, i.e., Λ_*pop*_ < 0 ([Sompolinsky et al., 1988]).

### 6.3 Recurrent Autoencoder setting

To analyze the dynamics of continuous attractor networks we first consider the dynamics of an auxiliary setting, which we call Recurrent Autoencoder (RAE) (Fig. S3A):

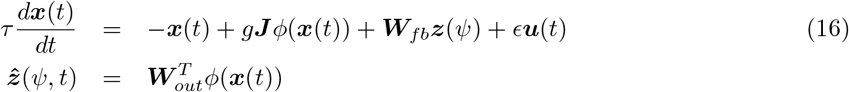

Here, we unfold the structured component of the recurrent connectivity from Eq.(1) (Fig.S 1Ci), and test what would be the *decoded* output of the RAE, 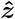, when externally enforcing its input through the *encoder, **W**_fb_*, to be ***z***.

The dynamics of the RNN (Eq.(1)) is governed by the discrepancy between the input and the output of the auxiliary system, 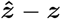. Namely, given a point ***x*** on the attractor, and its corresponding low-D projection ***z***, the dynamics of Eq.(1) should regenerate the same point persistently. In the RAE setting, this regeneration implies 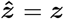. Conversely, a point ***x*** which is not on the attractor, would not be regenerated perfectly, resulting in a flow in the dynamics of Eq.(1).

As a result of the difference between dynamics along the on- and off-manifold directions, any neural trajectory in the vicinity of the manifold will first converge in the N-1 dimensions orthogonal to the 1D manifold (Fig.S 3D,G), and will then exhibit slower dynamics, approximately along the manifold itself (Fig.S 3G). Consequently, it is useful to consider the tangential error of the RAE, which governs the latter, slow, phase of the dynamics. This error is given by projecting the 2D error vector, 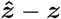, on the tangent to the manifold at point *ψ* and is approximated by:

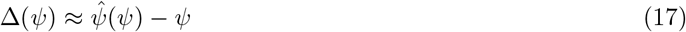

It measures the difference between the angle of the reconstructed point of the RAE, 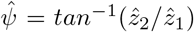 and the input angle *ψ* (Fig.S 3A,G).

### 6.4 The gain of the RAE

Linearizing Eq.(16) around a putative fixed point, ***x***(*ψ*), yields:

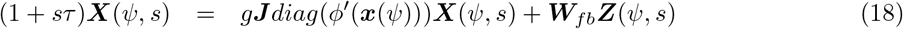

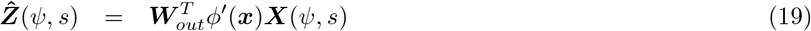

where ***X***(*ψ, s*) is the Laplace transform of the linearized state. The gain of the (open loop) RAE is defined by:

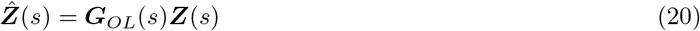

and can be computed as:

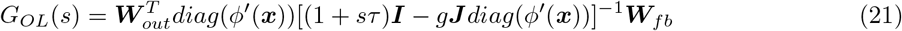

By closing the loop, i.e. by setting 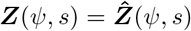 in Eq. (18), we obtain the gain of the fully recurrent network:

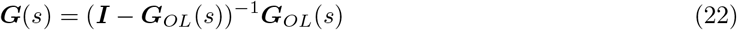

***G***(*s*) is a 2 × 2 matrix of *N* degree polynomial ratios and the poles of its determinant correspond to a subset of eigenvalues of the stability matrix of Eq.(14). In the large *N* limit, this degree is vastly reduced as the majority of linear dynamical modes become not observable via readout ***Z***, and it is only a small number of modes that persists in (20) and hence in (22) [Rivkind and Barak, 2017]. Specifically, these are the eigenvalues that appear due to the structured component of the connectivity and the bulk of remaining eigenvalues are not affected and obey a circular law (Fig. 5A and Fig. 7).

Marginal stability emerges when det(***G***) has a pole at *s* = 0. Equivalently, it follows from Eq.(22) that ***G**_OL_*(*s* = 0) must have an eigenvalue one, with the eigenvector being the tangent vector, 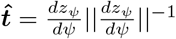. To further analyze the dynamics it is convenient to consider the gain of the RAE in the coordinates of the tangent and normal directions, with the normal direction, 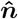, defined by a clockwise rotation of 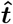 by *π*/2. We define the gain in the tangent direction to the manifold as

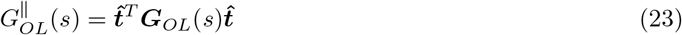

and, similarly, the gain in the normal direction as 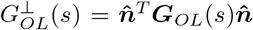. The conditions for a stable manifold are then that 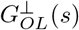 obeys stability conditions of a scalar feedback system, (e.g. Nyquist criterion [Nyquist, 1932]), while 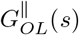 is required to obey marginality:

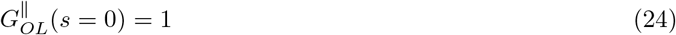

In the sequel, we argue that local dynamics along the manifold in this case can be approximated by first order differential equation, even for large *g* (see Section 6.7.3). Consequently, it must have the form:

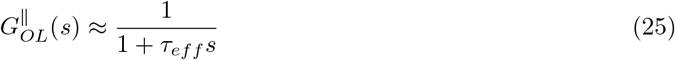

Finally, we note that small and slow translations along the manifold do not induce displacement at the normal direction, implying 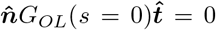. The second cross term 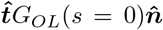 is nonvanishing in a general case, however, by symmetry considerations it vanishes if the manifold features reflection symmetry around 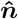. In particular, this condition holds at any point for a ring geometry, and for reflection points in the parameterized manifolds of Fig. 6, 7 and Fig.S 7.

### 6.5 Drift and connection to RAE framework

We assume that the dynamics in the N-1 directions orthogonal to the 1D manifold, 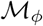, is convergent, and that the tangential error following training is small (or, alternatively, *ϵ* ≪ 1 in the case of external input). In this case, translations on the manifold are slower than the dynamics in the N-1 orthogonal directions. Under this assumption of timescale separation, we can link the drift along the manifold to the tangential error, as shown in the main text (Eq.(4)).

We start by considering the linearized RAE in the transnational direction. Adding a constant error Δ(*ψ, t*) = Δ(*ψ*) yields:

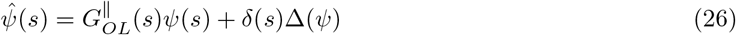

where *δ* is Dirac delta function. Closing loop by setting 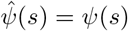 implies:

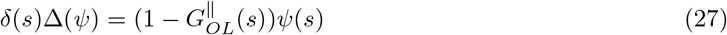

and assuming we can write the gain in the transnational direction as in Eq.(25), we get:

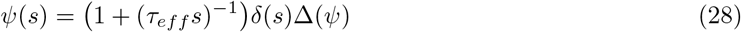

Recalling that the factor *s*^−1^ in Laplace domain translates into integral in the time domain yields:

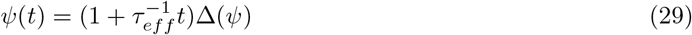

The velocity of drift along the manifold then follows:

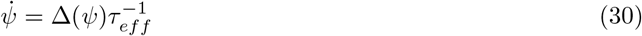

recapitulating Eq.(4) in the main text. Linerazing Eq.(30) around a fixed point:

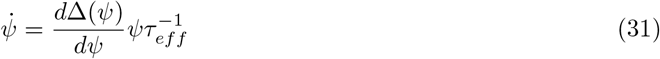

and:

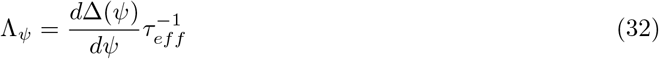

with stability achieved for Λ_*ψ*_ < 0, and marginality for Λ_*ψ*_ = 0. Here, we assumed (without loss of generality) that the fixed point is located at *ψ* = 0.

### 6.6 Trained manifold attractors

We train the network by sampling *M* ≪ *N* points of a pre-defined manifold, 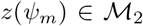, with *m* = 0…*M* – 1, 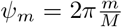 and run the dynamics in an open loop RAE setting (Eq.(16), Fig. S3) with *ϵ* = 0 until the recurrent dynamics converges [Jaeger, 2001, Sussillo and Abbott, 2009]. We obtain *M* states {*ϕ*(*x_m_*)}_0≤*m*≤*M*−1_ and train the decoder by minimizing the reconstruction error:

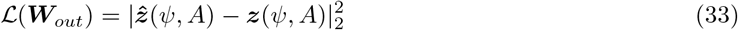

with 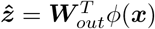. The least square (LS) solution yields:

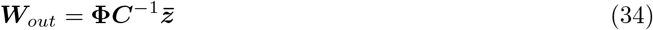

where here 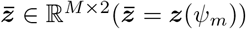, the steady state solutions of Eq.(16) are

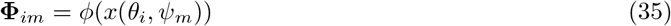

and the correlation between the rates is:

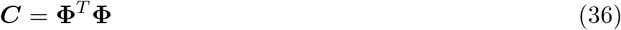

A point on the manifold can thus be written using the correlation matrix:

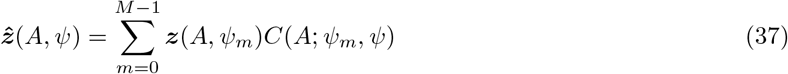

We next consider the singular value decomposition (SVD) of **Φ**:

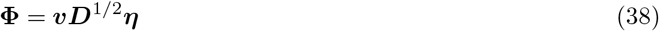

where 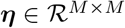 and 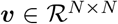 are the right and left singular vectors. The matrix 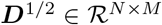 is the singular value (SV) matrix with 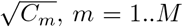, being the SVs and *C_m_* being the elements of the spectrum of the correlation matrix of Eq.(36). The decoder is thus given by:

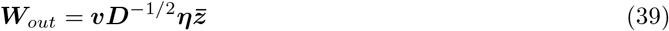

Due to stability requirements (Fig. 5A and Section 6.2), the above training procedure does not guarantee the emergence of a manifold attractor. First, a translational mode with marginal stability needs to emerge from sampling only a finite set of M points (red arrow in Fig. 5A). Second, amplitude direction needs to be stabilized (blue arrow in Fig. 5A). Finally, spontaneous activity must be suppressed [Rajan et al., 2010, Mastrogiuseppe and Ostojic, 2018] (purple arrow in Fig. 5A). To assess that, we developed a mean-field approach, calculated the steady state mean-field solution of Eq.(11), and analyzed its stability.

### 6.7 Trained ring attractor model in network with heterogeneous connectome

To train a ring manifold on top of heterogeneous connectivity (*g* > 0 in Eq. (1)) we apply the least square learning rule from Eq. (34) to *M* samples *z*_0..*M*-1_ from the ring curve 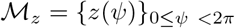 with:

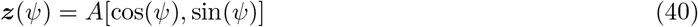

We will argue in the sequel (Eq. (46)) that in the limit of large *N* and for any *M* the correlation matrix is circulant. Consequently ***z***(*ψ*) is an eigenvector of ***C*** and the least square solution (39) is of a particularly simple form:

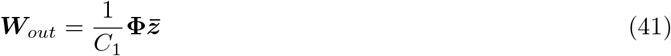

Remarkably, as depicted in Fig. 1E and as quantified by Eq.(54), circulant property of correlation matrix does not imply symmetry between individual neuronal representations. Consequently, the structured part of the recurrent interactions, ***W***, is not symmetric any more: while feedback weights are assumed to be 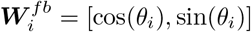 as in the symmetric-connectome model, this is not the case for the learned readout weights: 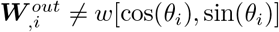 for any factor *w*.

With readout vector defined in terms of population activity, we are ready to develop a mean-field estimate for the open loop RAE gain (20).

#### 6.7.1 Mean-field solution for the neural representation of the manifold

To obtain the mean-field solution in real space we decompose the steady state solution of Eq.(16) into its deterministic and stochastic parts [Rajan et al., 2010, Rivkind and Barak, 2017]:

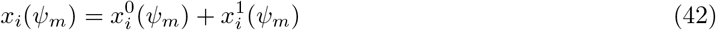

where the deterministic part, independent of disordered connectivity *J*, is

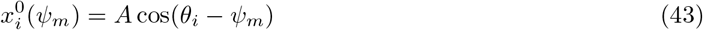

and the stochastic part, also known as the quenched disorder, is given by:

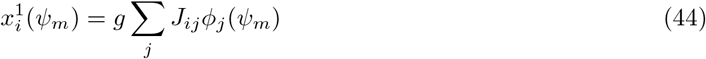

In the large *N* limit, 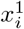 is replaced by a Gaussian random variable, *σy*, with *y* being a zero mean and unity variance Gaussian r.v. The variance, *σ*^2^ needs to be evaluated self-consistently. This is done by projecting (44) on itself:

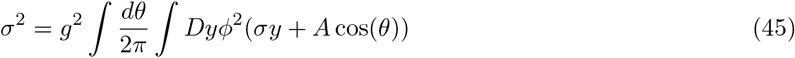

with 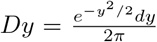. Importantly, due to the rotational symmetry in the deterministic part of Eq.(43), *σ* is independent of *θ* and *ψ_m_* in the large *N* limit. Moreover, since we assumed that *φ*(–*x*) = –*φ*(*x*), there is no bias term in Eq.(45).

Correlations between the inputs to the neurons on the manifold, *c*(*ψ_m_, ψ_m′_*) = 〈*x_i_*(*ψ_m_*)*x_i_*(*ψ_m′_*)〉, can be computed self-consistently as well. This is done by projecting Eq.(44) for *ψ_m_* onto the same equation but with *ψ_m′_*. In the specific case of a ring geometry, the latter only depends on the difference *ψ_m_* – *ψ_m′_*. To simplify notations, we assume *ψ_m′_* = 0 and obtain:

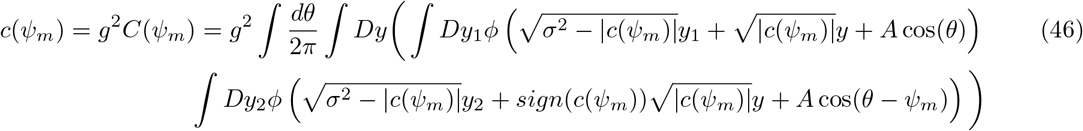

such that *c*(0) = *σ*^2^, and where we denote the correlations among the rates of the neural state by capital *C*:

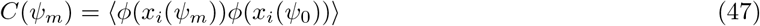

#### 6.7.2 Representation in Fourier space

Instead of mean-field estimate in the real space, we can also write the steady state solution of the dynamics using the singular value decomposition of Eq.(38). This turns out to be handy when considering the stability analysis, as we can write the decoder using the SVD. Using Eq.(38), the quenched disorder is

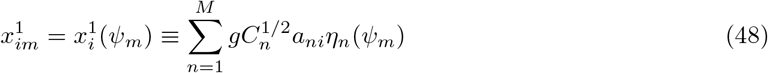

where 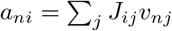 are Gaussian r.v. with zero mean and unit variance (and we assume 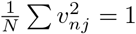).

In the case of the ring attractor, the correlation matrix is circulant (Fig. 2A). As a result, the SVs are simply the spatial Fourier modes, and the spectrum of the correlation function is:

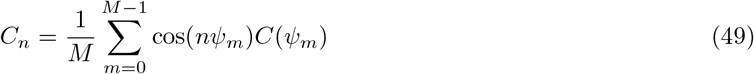

In this case, we denote the coefficients of even (resp. odd) modes corresponding to cos (resp. sin) functions by *a* (resp. *b*), as opposed to a case of general principal component decomposition where we do not make such a distinction (see Section 6.8). Eq.(48) thus yields:

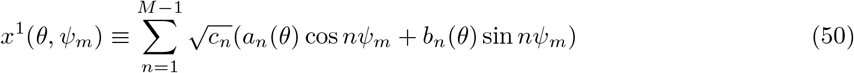

where we took the limit of *N* → ∞ such that *a_ni_, b_ni_* = *a_n_*(*θ*), *b_n_*(*θ*) and where we define

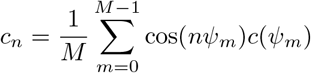

We can thus write the steady state solution using the Gaussian r.v. *a_n_, b_n_*:

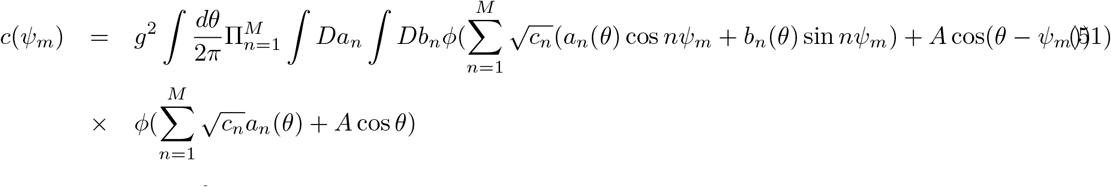

and as before, with *σ*^2^ = *c*(0). Thus, the statistics of the neuronal activity is fully specified in large *N* limit. In principle, to span the correlation *c*(*ψ_m_*) Eq.(51), one would need to integrate over a large number, 2*M*, of random variables *a_n_, b_n_*. However, as the spectrum of the correlation function decreases rapidly when *n* is increased, the amplitude of the higher frequency components in the quenched disorder is increasingly small. We thus write the approximate MF soultion in Fourier space by taking the approximation

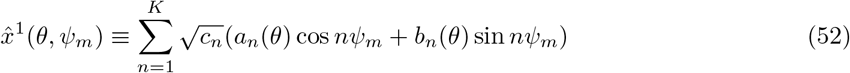

and where we defined the cut-off frequency, *K* (*K* < *M*).

At this point, we can also estimate the diversity of tuning curves [Cremers and Klugkist, 2018]. The tuning curve of the *i^th^* neuron is a function of the angle, *ψ*:

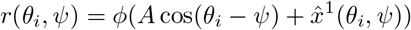

We can thus use the Gaussian statistics of *x*^1^(*θ_i_, ψ*) to generate tuning curves with the same statistics as in simulations. We define the selectivity index of a neuron *j* as:

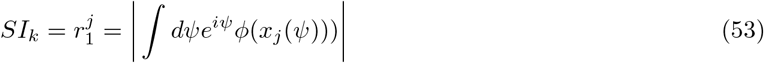

and we can further calculate the SD of the selectivity index, yielding:

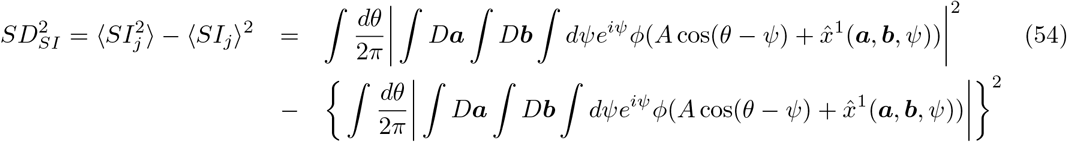

and ***a**, **b*** refers to product of all the coordinates up to the order *K*. Here again, in principle the calculation would require 2M random variables per one neuron, but in practice we find that *K* = 5 is accurate enough.

#### 6.7.3 Dynamics in the vicinity of the attractor - mean-field solution

Equipped with the mean-field solution for the neuronal state on the attractor, we are now set to explore the dynamics in its vicinity by evaluating Eq.(20) and (22). Rather than applying Eq.(21) directly, we estimate the elements 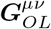 of 2×2 matrix ***G**_OL_* by the mean-field approximations of 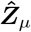 (Eq.(19)) driven by the appropriate input ***Z_ν_***(*s*) = ***ê_ν_*** (with ***ê_ν_*** a unity vector id the direction *ν*). Recalling that the linear decoder ***W**_out_* is spanned by rate-states *ϕ*(*x_i_*(*ψ_m_*)) (Eq.(34)):

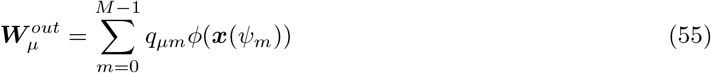

with *ν, μ* ∈ {1, 2} denote directions in the decoder plane, the terms of the RAE gain matrix (20) are:

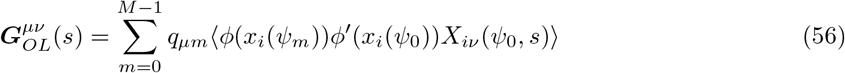

Similarly to Eq.(42), we decompose the linearized response:

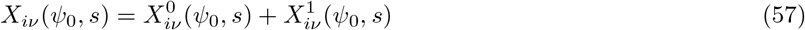

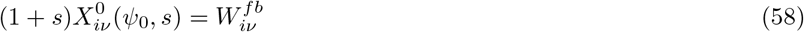

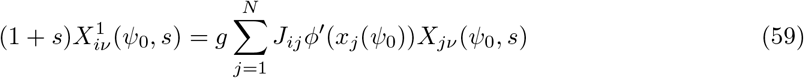

Multiplying Eq.(59) by Eq.(44) and projecting on the right singular vectors, *η_nm_*, we obtain:

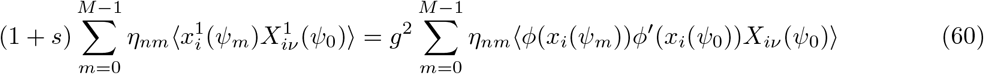

for which the RHS of Eq.(60) is closely related to the gain of the RAE (Eq. (56)).

Due to circulant property of ***C*** in the case of a ring geometry, the Fourier modes are the principal components of **Φ**, and *q_lm_* takes a particularly simple form (Eq.(41)): 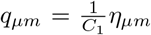 (*μ* ∈ {1, 2}). It is thus convenient to solve the problem in the Fourier domain with respect to *ψ*. For the sake of convenience, we focus on the point *ψ* = 0. We express *X* in the corresponding basis of quenched disorder according to (50):

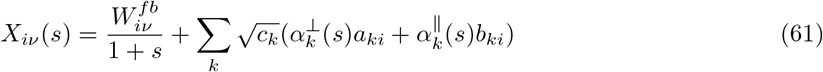

where we omit the argument *ψ*_0_ to simplify notations and *α*^⊥^(*s*) (resp. *α*^║^(*s*)) are coordinates of *X*(*s*) in the basis of {*a_k_, b_k_*}. Here, *μ, ν* = 1 correspond to the direction normal to the manifold and *μ, ν* = 2 corresponds to the tangential direction. We start by analysing the tangential direction, and we use the symbol ║ as shorthand for indexes 2 and 2, 2 (and similarly ⊥, for 1 and 1,1). Multiplying Eq.(50) with Eq.(61), and using Eq.(60), yields a self-consistent equations for the projections of linearized perturbation on the tangential direction of the manifold:

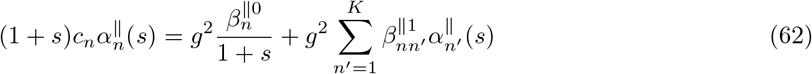

with

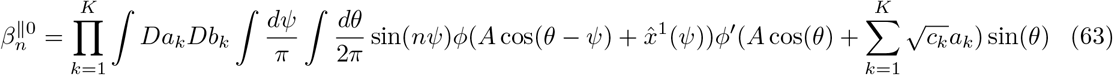

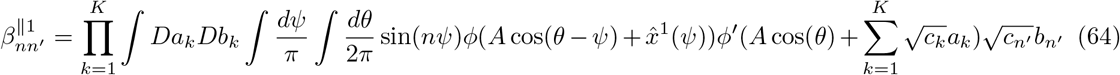

where we took the limit *M* → ∞ (assuming *M* ≪ *N*) and 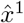 is an approximation of Eq.(50), up to *K*-th order (see Eq.(52)).

To obtain the gain of the RAE we note that 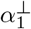 is closely related to the tangential gain. Indeed, from Eq.(62) and (60) the gain of the autoencoder in the manifold direction now follows:

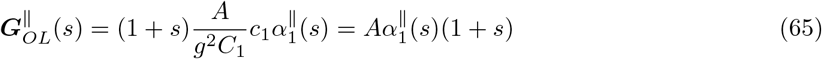

and 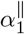 is the first element of vector *α*^║^:

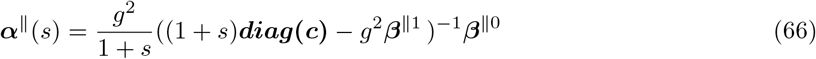

How many terms *K* are needed to estimate *α*^║^(*s*) and hence 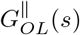? Using Stein’s lemma, we write Eq.(64) as

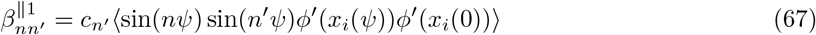

This term is small, unless *n* = *n′* = 1 and numerical simulations also show that first order approximation *K* = 1 is accurate for the relevant range of *s* (not shown).

We can now recover the effective timescale *τ_eff_* that governs the dynamics along the manifold (Eq.(4)). By assuming first order ansatz for the gain of the RAE we get:

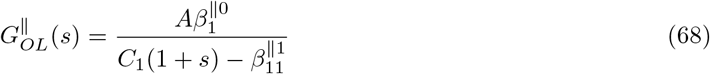

Reorganizing the above equation and recalling that the gain of the RAE at zero frequency is unity (see Eq.(24), yielding 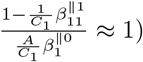 gives:

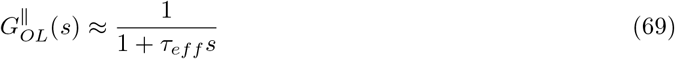

with the effective timescale given by:

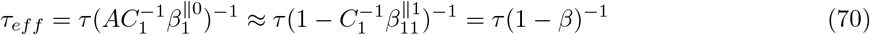

and where using Eq.(67) we get Eq.(5) in the main text:

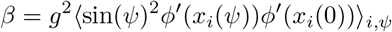

Following the same analysis, similar equations control the gain in the amplitude direction, but with:

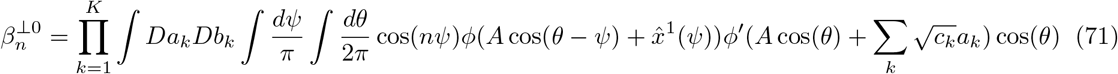

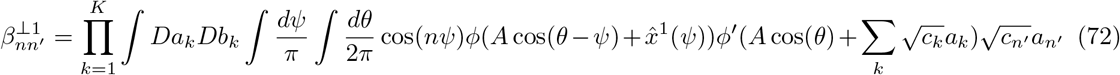

the gain 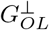 is computed similarly to Equations (65), (66) with leading eigenvalue Λ_*ρ*_ of the full system corresponding to the leading pole of the closed loop gain 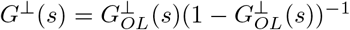.

The form of the terms 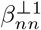 is more involved than in the case of tangential direction. Here, by using Stein lemma we get

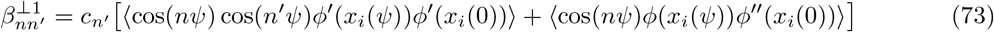

for which decay of high order terms is less obvious than in (67). Still, numerical calculations of 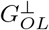 suggest that it also follows a first order dynamics (not shown).

#### 6.7.4 Effect of learning a finite number of samples from the manifold

When the number of trained points, *M*, is smaller than the number of neurons, *N*, the error is zero at these points. However, as long as *M* is finite the derivative of the tangential error Δ′ is not zero, implying that there is no marginal stability along the manifold (Eq. 31). Nevertheless, we now show that this derivative becomes small as the number of points grows, and marginal stability is being effectively recovered.

For a ring geometry, and in the large *N* limit, the sampled points are equivalent and, without loss of generality, we calculate the derivative of the tangential error, Δ′ (31) at *ψ*_0_ = 0. The tangential error at *ψ*_0_ = 0 is given by 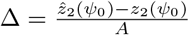. Substituting into (41) we obtain:

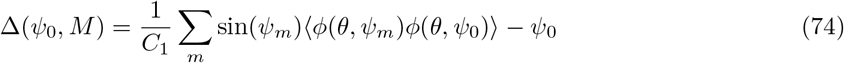

yielding:

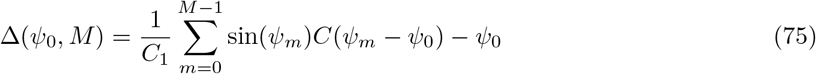

To obtain Δ′ we define the continuous correlation function, *C*(*ψ*). In this sense, the notation *C′*(*ψ_m_*) refers to 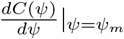. We expand the (continuous) correlation function, which is an even function of *ψ*, in Fourier series as 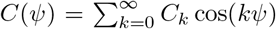 and write the derivative Δ′ of the tangential error at the sampled points:

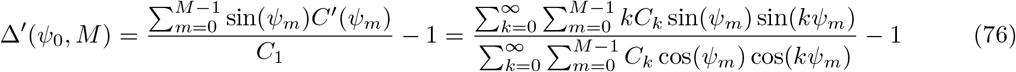

First, it is now obvious from RHS of (76) that lim_*M*→∞_ Δ′ = 0, implying that manifold direction exhibits marginal stability in the case of infinitely many training samples. Furthermore, correction due to finite *M* is expressible via spectrum of *C*. Namely:

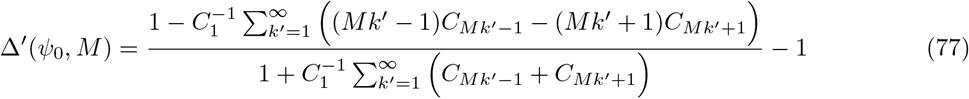

In case of a fast decaying spectrum of the correlation function, such that *C_k_* ≫ *C_k′_* for *k′* > *k*, as in cases shown in Fig. 4D, Eq.(77) can be approximated by:

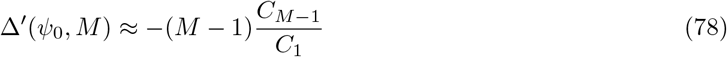

We can now plug this result, along with the effective time constant from (70) into (32) to obtain mean-field estimate for the maximal eigenvalue in the tangential direction:

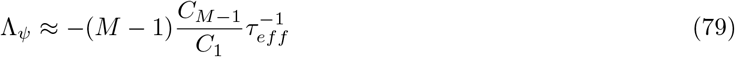

We thus conclude that marginally stable direction, which enables a continuum of persistent states, is recovered exponentially fast with the number of samples, *M* (exponential rate is due to the exponential decay of Fourier coefficients of correlation function, which is true for infinitely differentiable correlation functions, see e.g. [Katznelson, 2004]).

### 6.8 General 1D manifold attractor

Similarly to the case of the ring manifold, we write the steady state mean-field solution and stability equations of the dynamics in Eq.(11) for a general 1D manifold. In contrast to the case of a ring manifold, in manifolds with general geometry the correlation matrix is no longer circulant and therefore the eigenvectors are not the Fourier modes. Still, similar analysis is applicable and similar mean-field equations control the stability.

For a general curve 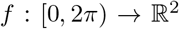 and ***z***(*ψ*) = ***f***(*ψ*), we follow Section 6.7.2 and write the correlation function in the singular vectors domain:

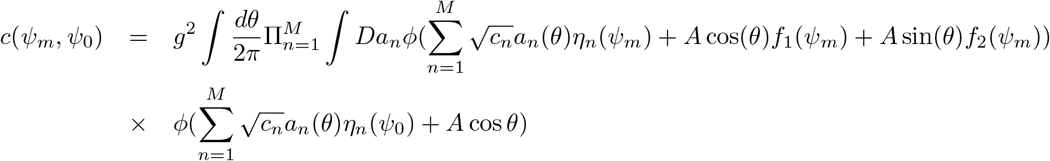

Next, stability is evaluated based on Eq.(60), with the only difference that the decoder is spanned by more than just two singular vectors. Recall that

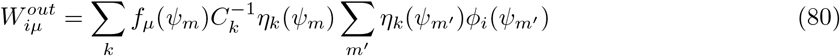

So that

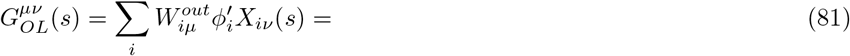

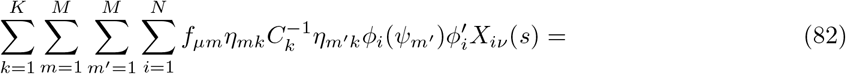

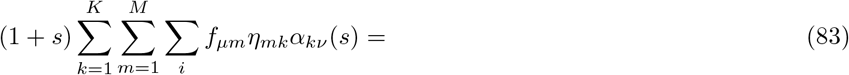

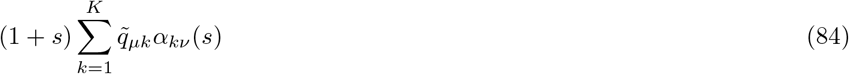

where we defined

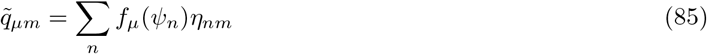

Following the derivation of Section 6.7.3, with

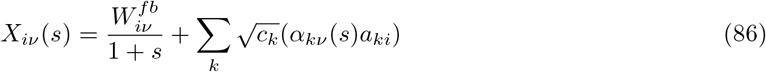

we reach a similar self-consistent equation for the order parameters 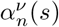 as in Eq.(62), but with

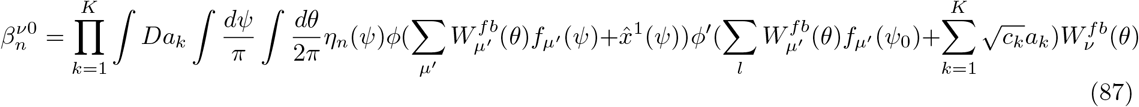

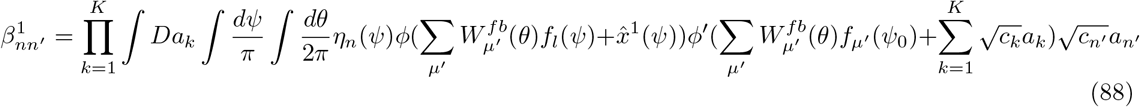

with *ψ*_0_ denoting the point at the manifold for which the gain is computed. Here, once again, we define the cut-off *K* < *M*. Note that the coefficients 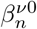 depend on the input index *ν*, while coefficients 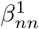 do not.

A notable difference from the ring geometry, where up to rotation transformation the gain matrix is the same for all points on the manifold, is that for a general geometry the gain varies qualitatively with *ψ*. Destabilization may occur for some values of *ψ*, while other regions remain stable. Another difference from a ring geometry is that for manifolds with large deformations, taking a cutoff at *K* = 2 (i.e. one mode per tangent direction and one mode for normal direction) is not enough and the anzats of Eq.(25) does not longer hold (Fig. 6–7). Indeed, the decoder is now spanned by a mixture of several singular vectors, and consequently more order parameters *α_nν_*(*s*) are needed to describe the dynamics.

Similarly to (61), the *α_n_*(*s*) in a case of general geometry are defined by:

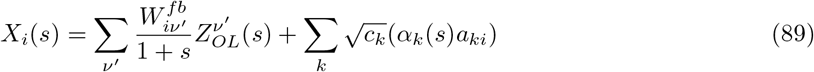

Specifically, given our system with input *Z_OL_*(*s*), rather that with *unity* input that was assumed in an open loop system, we have *α* given by:

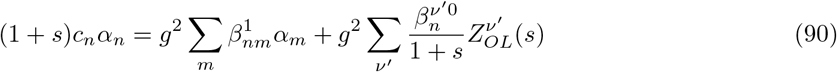

and the readout 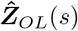 can be obtained from (84) as:

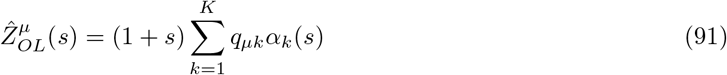

closing loop is done by imposing 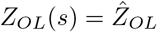 which turns (84) into:

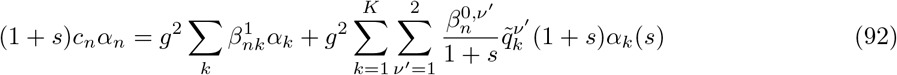

this can be re-written in a form:

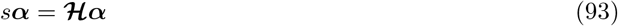

where the *K* × *K* mean-field stability matrix is:

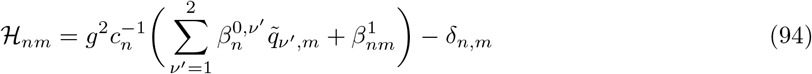

*K* eigenvalues of the mean-field stability matrices 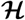 correspond to the change in *K* eigenvalues of the spectrum of the stability matrix ***H*** (Eq.(14)) as a result of adding the trained structured matrix ***W*** to the random matrix *g**J***.

In points with reflection symmetry in the target function ***f***, further simplification is possible. The matrix 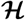 in such cases is decomposable into blocks of odd and even principal components with no interaction in between them. The block of even principal components 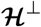 will describe the stability in the normal (amplitude) direction while the block of odd modes 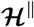 will account for the tangential direction. Obviously, this is the case for perfect ring geometry, for which such an even-odd dissection presented in Eq. (61) and the even and odd modes are given by cos(*kψ*) and sin(*kψ*) respectively. Furthermore, this is also the case for the minima and maxima of amplitude in the manifolds that were analysed here (Fig. 7).

### 6.9 Symmetric-connectome ring model – revisit in RAE setting

For completeness, we apply the RAE analysis for the classical symmetric-connectome ring attractor model [Ben-Yishai et al., 1995, Mastrogiuseppe and Ostojic, 2018]. This case corresponds in our framework to setting feedback and output weights to be: 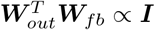 and setting *g* = 0. Thus, when we unfold the structured component of the connectivity the RAE (Eq.(16)) becomes a simple feedforward autoencoder. The strength of the structured recurrent loop is determined by 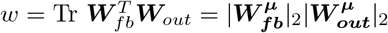 with *μ* = 1, 2 and it equals *w* = *J*_2_ in the original notation of [Ben-Yishai et al., 1995].

Following [Ben-Yishai et al., 1995], one example for such a choice is 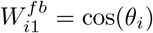 and 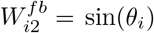, with *θ_i_* = 2*πi*/*N* (see [Mastrogiuseppe and Ostojic, 2018] for a different choice of orthogonal vectors). The decoder is then normalized such that 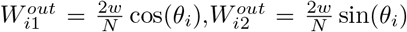, yielding the cosine connectivity profile: 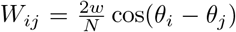. The parameter w thus controls the amplitude of the bump through the self-consistent equation:

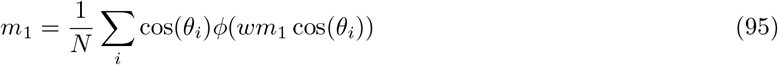

To follow the recurrent autoencoder framework, the representation in 2D plane of the RAE needs to be obtained. Specifically, we need to compute *A* of Eq.(12). On the one hand, since every point on the manifold attractor is a fixed point, we get that for *ψ* = 0 (as well as for any other 0 ≤ *ψ* ≤ 2*π*):

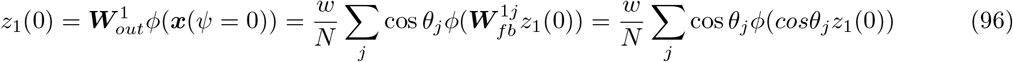

which together with Eq.(95) yields *z*_1_(0) = *wm*_1_. On the other hand, according to notation of Eq.(12) *z*_1_(*ψ* = 0) = *A*. Consequently:

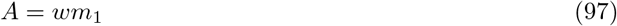

We are now set to evaluate local dynamics and, in particular, the drift velocity: It follows from (18), that for unity input in the t direction *Z*(*s*) = (0; 1) linearized state is given by

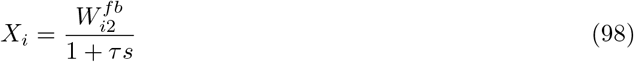

and the open loop gain in the tangential direction is:

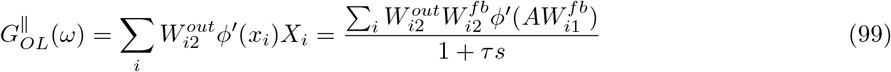

Approximating sum by integral, and integrating by parts we obtain:

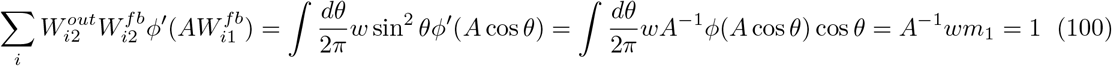

that is, we recovered the general formula (25): albeit with *τ_eff_* = *τ*.

#### 6.9.1 Input response

After obtaining the gain of the autodecoder, we proceed with the response of the symmetric-connectome network to external inputs. We assume that ***W**_in_* = ***W**_fb_* and that input of a strength *ϵ* is introduced at direction *ψ*_1_ to a system located at *ψ*_0_ = 0. We then have Δ*z* ≈ *ϵ*(cos(*ψ*_1_) sin(*ψ*_1_)). In case of *ψ*_0_ =0 we have *z*_0_ = (*A*, 0) and the tangential RE is 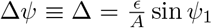. By the virtue of (30) the rotation speed is:

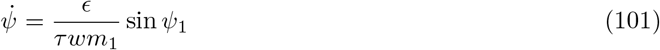

in agreement with the original result by [Ben-Yishai et al., 1995

#### 6.9.2 Effect of adding noise

For a structured connectivity, the amplitude *m*_1_ can be computed self-consistently as:

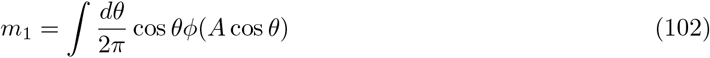

with *A* defined by (97). With synaptic heterogeneity in presence, *m*_1_ needs to be evaluated self-consistently together with the variance of the quenched disorder:

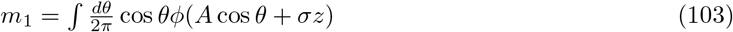

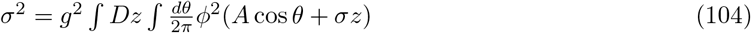

To estimate the mean-squared drift velocity we use equation (30) and note that in case of small *g* we have *τ_eff_* ≈ *τ*. It remains to estimate the mean-squared error. This can be done by explicitly by computing the variance of the readout:

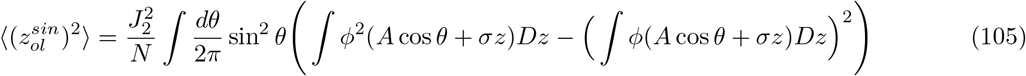

In case of small *g* – and hence small *σ* – we can linearize the above variance and use:

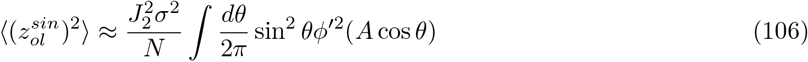

To obtain Δ, we normalize the squared error, *z*, by the amplitude, *A*, and get:

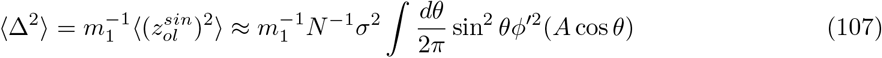

which can be compared to [Itskov et al., 2011] and which is compared with simulations in Fig. S1E.

## Notes

### Competing Interest Statement

The authors have declared no competing interest.

### Summary of Updates

Improved last part of results to clarify dimensionality reduction. Improved methods. Fixed typos.

